# Infection characteristics among *Serratia marcescens* capsule lineages

**DOI:** 10.1101/2024.08.23.609398

**Authors:** Mark T. Anderson, Stephanie D. Himpsl, Leandra G. Kingsley, Sara N. Smith, Michael A. Bachman, Harry L. T. Mobley

## Abstract

*Serratia marcescens* is a healthcare-associated pathogen that can cause severe infections including bacteremia and pneumonia. The capsule polysaccharide of *S. marcescens* is a bacteremia fitness determinant and previous work defined capsule locus (KL) diversity within the species. Strains belonging to KL1 and KL2 capsule clades produce sialylated polysaccharides and represent the largest subpopulation of isolates from clinical origin. In this study, the contribution of these and other *S. marcescens* capsules to infection was determined in animal and cellular models. Using a murine model of primary bacteremia, clinical isolates of multiple KL types demonstrated capsule-dependent colonization of spleen, liver, and kidney following tail vein inoculation. Similar results were observed using a bacteremic pneumonia model, in that all tested strains of clinical origin demonstrated a requirement for capsule in both the primary lung infection site and for bloodstream dissemination to secondary organs. Finally, capsule from each KL clade was examined for the ability to resist internalization by bone marrow-derived macrophages. Only the sialylated KL1 and KL2 clade strains exhibited capsule-dependent inhibition of internalization, including KL2 capsule produced in a heterologous background. Together these findings indicate that lineage-specific resistance to macrophage phagocytosis may enhance survival and antibacterial defenses of clinically-adapted *S. marcescens*.

**IMPORTANCE:** Bacteremia occurs when the host immune system fails to contain bacterial bloodstream replication following an initial inoculation event from either an internal or external source. Capsule polysaccharides play a protective role for *Serratia marcescens* during bacteremia but there is abundant genetic diversity at the capsule-encoding locus within the species. This study compares the infection characteristics of *S. marcescens* isolates belonging to five capsule types and defines the contributions to infection fitness for each. By characterizing the differences in capsule dependence and infection potential between *S. marcescens* strains, efforts to combat these life-threatening infections can be focused toward identifying strategies that target the most critical genetic lineages of this important opportunistic pathogen.

## INTRODUCTION

*Serratia marcescens* is one the common causes of bacteremia and pneumonia among Gram-negative bacterial species (1, 2) and an estimated 100,000 deaths (42,000 associated with drug resistance) were due to *Serratia* species in 2019 alone (3, 4). Many of these infections occur in individuals with pre-existing conditions or during prolonged hospital stays. Pediatric populations are also vulnerable to severe *S. marcescens* infections, often occurring as nosocomial outbreaks (5–7). In addition to the systemic and life-threatening infections that are the focus of this work, *S. marcescens* is capable of a wide range of other pathogenic interactions with both human and non-human hosts (8–11). The clinical significance of *S. marcescens* is contrasted by the prevalence of the species in many environments, with isolation sources ranging from soil, water, plants, and insects (8) and highlighting the range of niches in which the organism thrives.

Genomic studies investigating the population structure of *S. marcescens* have defined the species-level diversity and have identified distinct lineages within the species. Some lineages have stringent correlation with clinical sources, consistent with niche adaptation to the infection environment and supported by a discrete repertoire of accessory genomic elements enriched within these clades (12–15). Other lineages are conversely associated with non-clinical or environmental sources. The clinical lineages have a higher proportion of antimicrobial resistance genes and there is evidence for substantial propagation of drug-resistant clades over time and geographic location (12, 14, 16). The major infection-associated genotypes are now represented by hundreds of sequenced strains enabling experimental examination of phenotypes that are predicted to impact *S. marcescens* pathogenesis. Our own work has also demonstrated a distinction between clinical and environmental *S. marcescens* lineages strictly through comparison of the locus encoding capsule polysaccharide (CPS) (17).

The CPS for one *S. marcescens* capsule type was characterized as a critical fitness determinant during bloodstream infection (18). Like other encapsulated Enterobacterales species, the CPS of *S. marcescens* is encoded in a genomic locus that varies extensively between isolates (17). Our comparison of capsule loci (KL) from infection isolates determined that clades KL1 and KL2 were overrepresented among a cohort of >300 genomes and that strains from both clades produced sialylated CPS. Ketodeoxynonulonic acid (KDN) was the predominant sialic acid identified from KL1 and KL2 strains but a minor proportion of *N*-acetylneuraminic acid (Neu5Ac) was also detected. In addition to these predominant clinical capsule types, less abundant capsule clades from clinical or non-infection sources were also defined. In this work, we sought to determine the infection characteristics of *S. marcescens* strains representing five different capsule clades using both animal and cellular model systems.

## RESULTS

### Infectivity of *S. marcescens* isolates following bloodstream inoculation

Capsule was previously determined to be an important fitness factor based on experiments using a single *S. marcescens* bacteremia clinical isolate belonging to clade KL1 (17, 18). As an initial assessment of infection capability for isolates differentiated by capsule type, strains selected from clades KL1-KL5 (Table 1) were inoculated into the bloodstream of mice via tail vein injection (TVI) and bacterial survival was measured at 24 h. KL1 strain UMH9 stably colonized the spleen, liver, and kidneys in a manner consistent with previous results (19) (Fig. 1). The KL2-5 strains also colonized blood-filtering organs at 24 h, but significant variation in bacterial burdens was observed among the strains in both the spleen and liver (Fig. 1A and 1B), with KL3 bacteremia isolate UMH7 consistently achieving the highest density in both organs. The KL2 (gn773) and KL4 (UMH11) bacteremia isolates were similarly elevated in liver compared to KL1 and the *S. marcescens* type strain ATCC 13880 (KL5), a pond water isolate. Although variability in the kidneys was higher than the other organs (Fig. 1C), the KL2 and KL5 strains trended towards lower colonization levels, approaching the limit of detection. Thus, while all clinical and non-clinical *S. marcescens* strains were capable of bacteremia, significant organ-specific colonization differences are observed.

**Table 1.**
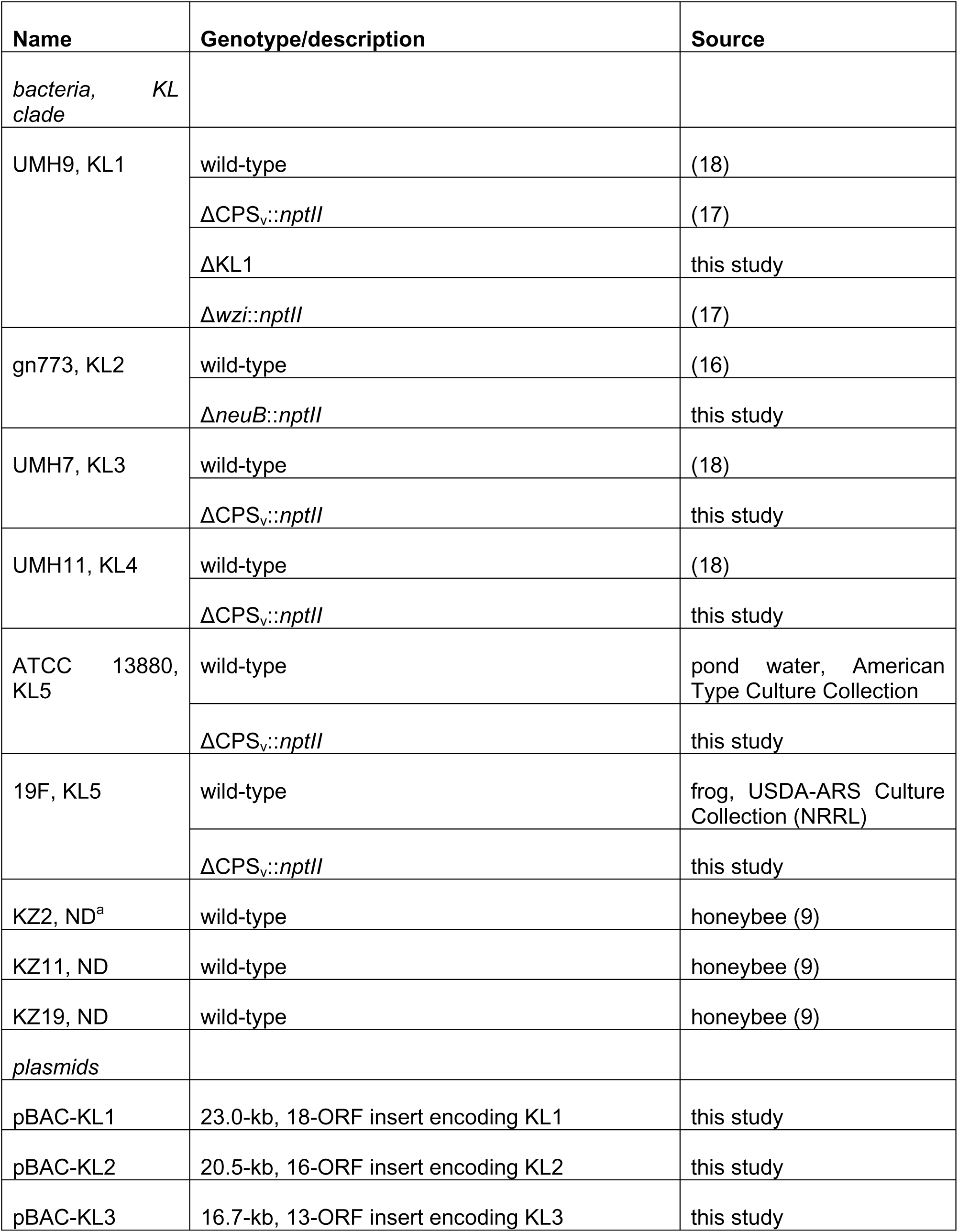
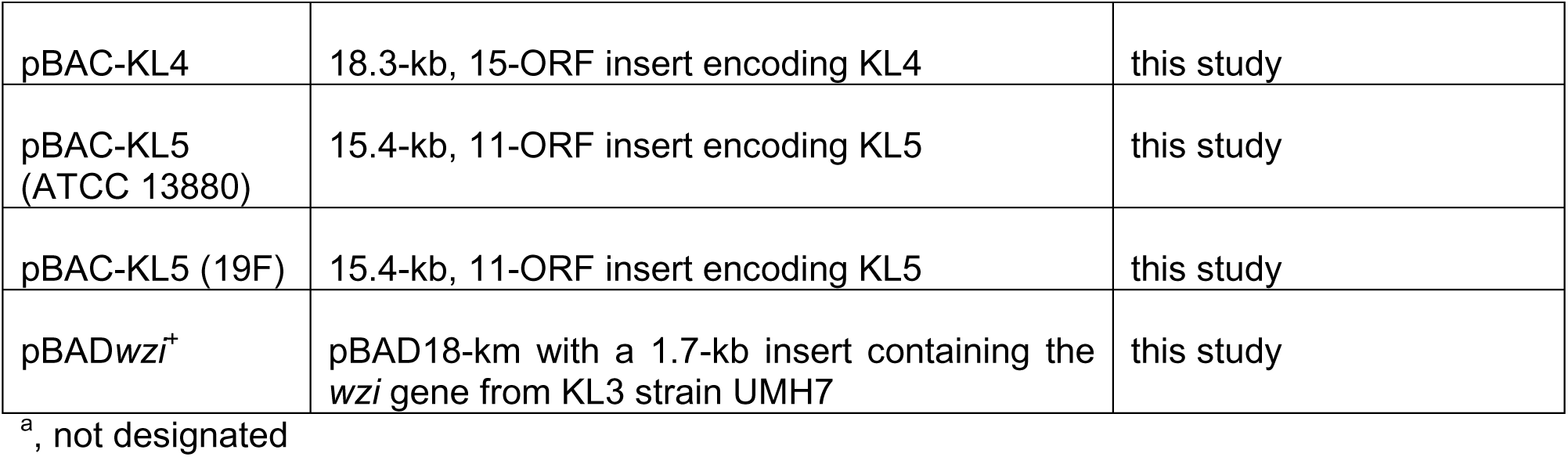
*S. marcescens* strains and recombinant CPS plasmids.

**Figure 1.**
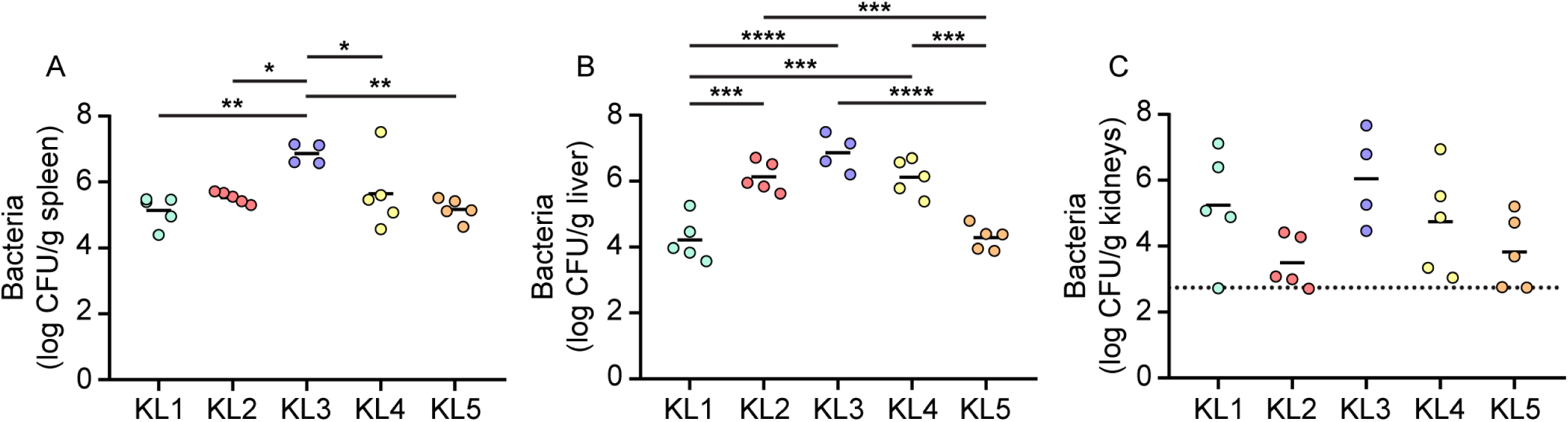
Strain variations in organ colonization following TVI bacteremia. *S. marcescens* strains were inoculated into C57BL/6J mice (n=5) via TVI and bacterial colonization in spleen (A), liver (B), and kidneys (C) was determined by viable counts. Log transformed mean bacterial burdens are indicated by the solid lines. Statistical significance was assessed by ordinary one-way ANOVA with Tukey’s multiple comparisons test: *, P < 0.05; **, P < 0.01; ***, P < 0.001; ****, P < 0.0001. The dotted line in panel C represents the highest value among samples that were at or below the limit of detection.

### Capsule contributions to bacterial survival following TVI

Acapsular mutants of each KL type were next generated to test the contribution of CPS to bacteremia across strains (Table 1). KL1, KL3, KL4, and KL5 strains were mutated such that the variable and clade specific region of each KL (CPS_v_) (Fig. S1A) (17) was deleted and replaced with an insert fragment encoding kanamycin resistance. Attempts to construct a similar ΔCPS_v_ mutation in the KL2 strain were unsuccessful, despite multiple efforts using different mutagenic systems. As an alternative, a capsule-null phenotype was achieved by disrupting the *neuB* gene encoding the sialic acid synthase within the KL2 CPS_v_ region, similar to a previously described capsule-null *neuB* mutant of KL1 (17). Initial assessments of these five mutants confirmed that the KL mutations disrupted high molecular weight CPS production, eliminated extracellular CPS uronic acids, and did not prevent O-antigen synthesis (Fig. S1). The only exception was KL5 ATCC 13880, which did not yield detectable CPS or O-antigen from either the wild-type or ΔCPS_v_ mutant.

The relative fitness of each capsule mutant was first quantitated in comparison to the wild-type parent strain in TVI mixed infections via competitive index (CI). All KL types under these mixed inoculum parameters exhibited similar total burdens in the spleen, kidneys, and liver of infected mice (Fig. S2). For each strain with detectable CPS production *in vitro* (Fig. S1), a significant competitive disadvantage in survival after 24 h was observed for mutants lacking capsular genes (Fig. 2A-D). The fitness advantage provided by KL4 CPS was only significant for bacteria in the liver but showed a similar trend in kidneys. Combined, these results demonstrate that capsule is important for bacterial survival across multiple clinical *S. marcescens* isolates and capsular clades. In contrast, the KL5 strain demonstrated no significant change in fitness upon CPS_v_ mutation (Fig. 2E). This result was anticipated given the lack of CPS associated with this strain; however, the six genes present in the ATCC 13880 KL5 CPS_v_ region (Fig. S1), all appear to be uninterrupted in the genome sequence (NZ_CP072199.1) and thus their functional significance is unclear. To further evaluate contributions of each CPS type to bloodstream fitness, strains were exposed to human serum for 90 minutes followed by enumeration of viable bacteria. KL1-KL4 CPS provided resistance to the bactericidal activity of serum that was at least 6-fold greater than acapsular mutant derivatives (Fig. 3A-D). No significant difference in serum susceptibility was observed between the KL5 strains (Fig. 3E).

**Figure 2.**
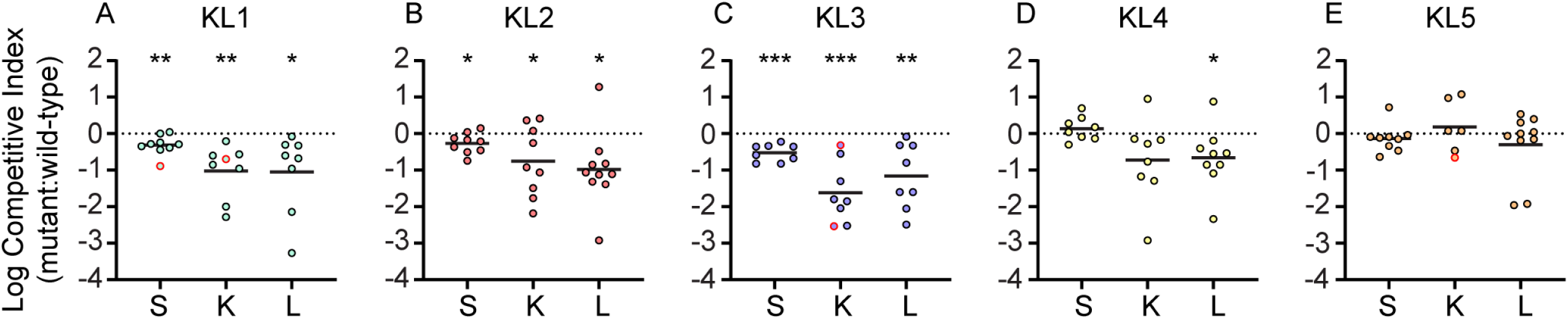
Requirement for capsule during TVI bacteremia. A-E. CI for bacteria recovered from 24 h mixed strain competition TVI infections in C57BL/6J mice (two cohorts of five mice, n=10). Solid lines represent the mean of log transformed values. Red outlined symbols indicate CI for which mutant bacteria were recovered at or below the limit of detection. Statistical significance was determined by one sample t-test with a hypothesized mean value of zero (dotted line), representing neutral fitness. Points below the dotted line represent samples in which the mutant strain was outcompeted by wild-type bacteria. Symbols and abbreviations: *, P < 0.05; **, P < 0.01; ***, P < 0.001. S, spleen; K, kidney; L, liver.

**Figure 3.**
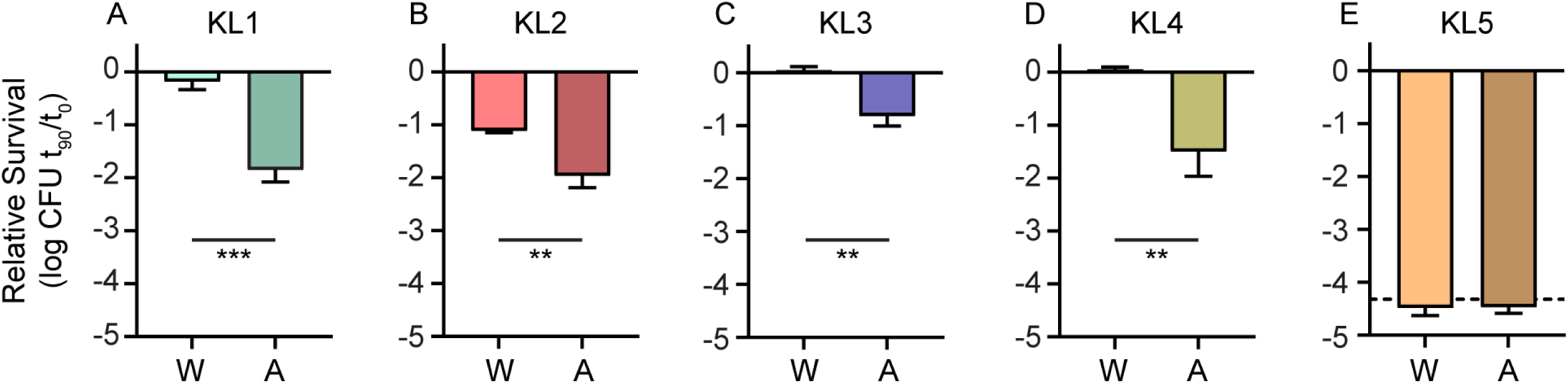
Capsules of clinical isolates protect from serum bactericidal activity. The survival of wild-type (WT) and acapsular (A) mutant strains was determined in the presence of 40% human serum after 90 minutes exposure relative to time zero with bars representing mean values (n=3). The dashed line indicates the limit of detection, where relevant. Statistical significance was assessed by Student’s t-test: **, P < 0.01; ***, P < 0.001.

### Capsule contributions to bacteremic pneumonia

Bacteremia frequently originates from localized infections that disseminate and become systemic (20), as opposed to primary bacteremia in which organisms gain direct access to the bloodstream via an exogenous source such as a hypodermic needle or intravenous catheter. A pneumonia model with secondary bacteremia was therefore developed based on a previously described *K. pneumoniae* model (21). Following retropharyngeal co-inoculation of wild-type and capsule mutant strains into mice, colonization of the lungs was observed at 24 h post-inoculation (Fig. 4A). The spleen, kidneys, and liver were also colonized at this time point (Fig. 4B-D) and to levels that approximated those observed in the primary bacteremia model (Fig. 1). *S. marcescens* escape from the lungs therefore occurs readily and results in stable organ colonization. The overall trends in bacterial burdens of the spleen, kidneys, and liver between this dissemination-dependent route and the TVI route were also similar for individual KL types in that KL3 bacteria exhibited the highest density followed by KL2 and KL4, then KL1 and KL5 (Fig. 1A-C and Fig. 4 B-D). Bacterial accumulation at systemic sites also correlated to primary lung burden when assayed at the time of sacrifice (Fig. 4A). Therefore, primary lung burden may influence dissemination kinetics and subsequent organ colonization in addition to any differences in infection capacity between strains.

**Figure 4.**
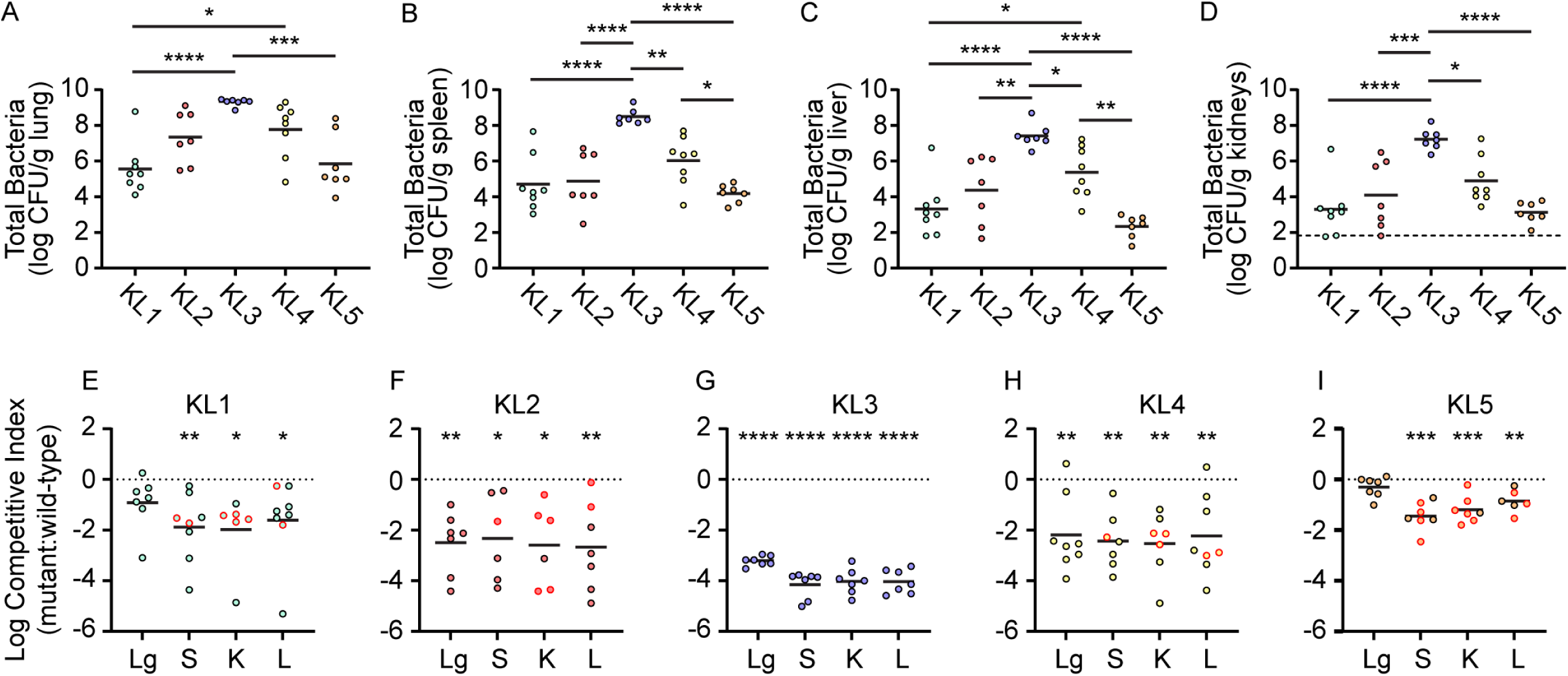
Requirement for capsule during bacteremic pneumonia. Combined wild-type and capsule mutant bacteria recovered from the lung (A), spleen (B), liver (C), and kidneys (D) of C57BL/6J mice (two cohorts of five mice, n=10) following mixed strain competition infections (24 h). Solid lines represent the mean of log transformed viable bacteria and the dotted line in panel D indicates the highest value among samples that were at or below the limit of detection from kidneys. Differences in bacterial burdens between strains were assessed by one-way ANOVA with Tukey’s multiple comparisons test: *, P < 0.05; **, P < 0.01; ***, P < 0.001; ****, P < 0.0001. E-I. CI comparing relative survival of capsule mutant and wild-type bacteria in lung (Lg), spleen (S), kidneys (K), and liver (L) for the infections shown in panels A-D. Symbols with red outlines denote CI from which capsule mutant strains were recovered at or below the limit of detection. Fitness defects were assessed by one-sample t-test against the hypothetical null value of zero (dotted lines) representing neutral fitness. Points below the dotted line represent samples in which the mutant strain was outcompeted by wild-type bacteria. Symbols: *, P < 0.05; **, P < 0.01; ***, P < 0.001; ****, P < 0.0001.

To define the requirement for capsule in pneumonia, the relative recovery of capsule mutant and wild-type bacteria for KL1-5 strains was determined. The four acapsular strains of clinical origin all demonstrated a severe competitive disadvantage compared to parental strains (Fig. 4E-H), with the mean recovery of capsule-deficient strains being *ca*. 100-fold or less than wild-type for most organ and strain combinations. Thus, in both primary and secondary bacteremia models, capsule is a critical fitness determinant across human-associated *S. marcescens*. Unexpectedly, a significant competitive disadvantage for the capsule mutant derivative of KL5 ATCC 13880 was also observed in the spleen, kidneys, and liver (Fig. 4I). Given the contrast between these results and those from TVI (Fig. 2E), the mutated KL5 capsular genes could play an active role in bacterial survival during lung dissemination. However, wild-type ATCC 13880 bacteria lacked a competitive advantage in the lung whereas the clinical strains demonstrated at least an 8-fold advantage in the lung compared to isogenic acapsular derivatives (Fig. 4E-H). Additional efforts to identify ATCC 13880 CPS by microscopy during repeated passage under selective pressure in human serum failed to provide evidence that CPS synthesis could be induced in this strain (Fig. S3).

### KL5 CPS does not contribute to primary bacteremia

The lack of KL5 CPS from ATCC 13880 was a limitation to understanding the role of CPS from non-clinical *S. marcescens*. To assess whether other KL5 strains produce CPS, a second representative designated 19F (Table 1) was acquired and characterized. High molecular weight polysaccharide was detected from 19F (Fig. 5A) and this strain had abundant surface-associated uronic acids (Fig. 5B), demonstrating that 19F synthesizes CPS. Three additional environmental strains that were originally isolated from honeybees (Table 1) also produce acidic CPS (Fig. S4), solidifying the conclusion that the acapsular phenotype of ATCC 13880 is not conserved among environmental strains. No significant fitness difference in TVI CI was observed between wild-type 19F and a ΔCPS_v_ mutant derivative (Fig. 5C), supporting the conclusion that the KL5 CPS contributes minimally to 19F bacteremia fitness. We hypothesized that the lack of a CPS-mediated fitness advantage for either ATCC 13880 or 19F would result in enhanced clearance of these strains compared to an encapsuled strain such as KL1 UMH9 and therefore survival following TVI was assessed over time. Both KL5 strains trended toward enhanced clearance compared to KL1, with ATCC 13880 showing significantly lower bacterial burdens than KL1 in all organs at 48 h post-inoculation and an overall loss of bacteria over time in the liver (Fig. 5D-F). Recovery of strain 19F was also lower than KL1 at 24 h in the spleen and kidney but did not show significant differences in the liver at any time point. Additionally, KL1 was the only strain capable of significant expansion over the course of the experiment, with high levels of bacteria found in the kidney by 48 h (Fig. 5E). Clearance of all strains appeared to be most effective in the spleen, since bacterial burdens decreased significantly over time (Fig. 5D).

**Figure 5.**
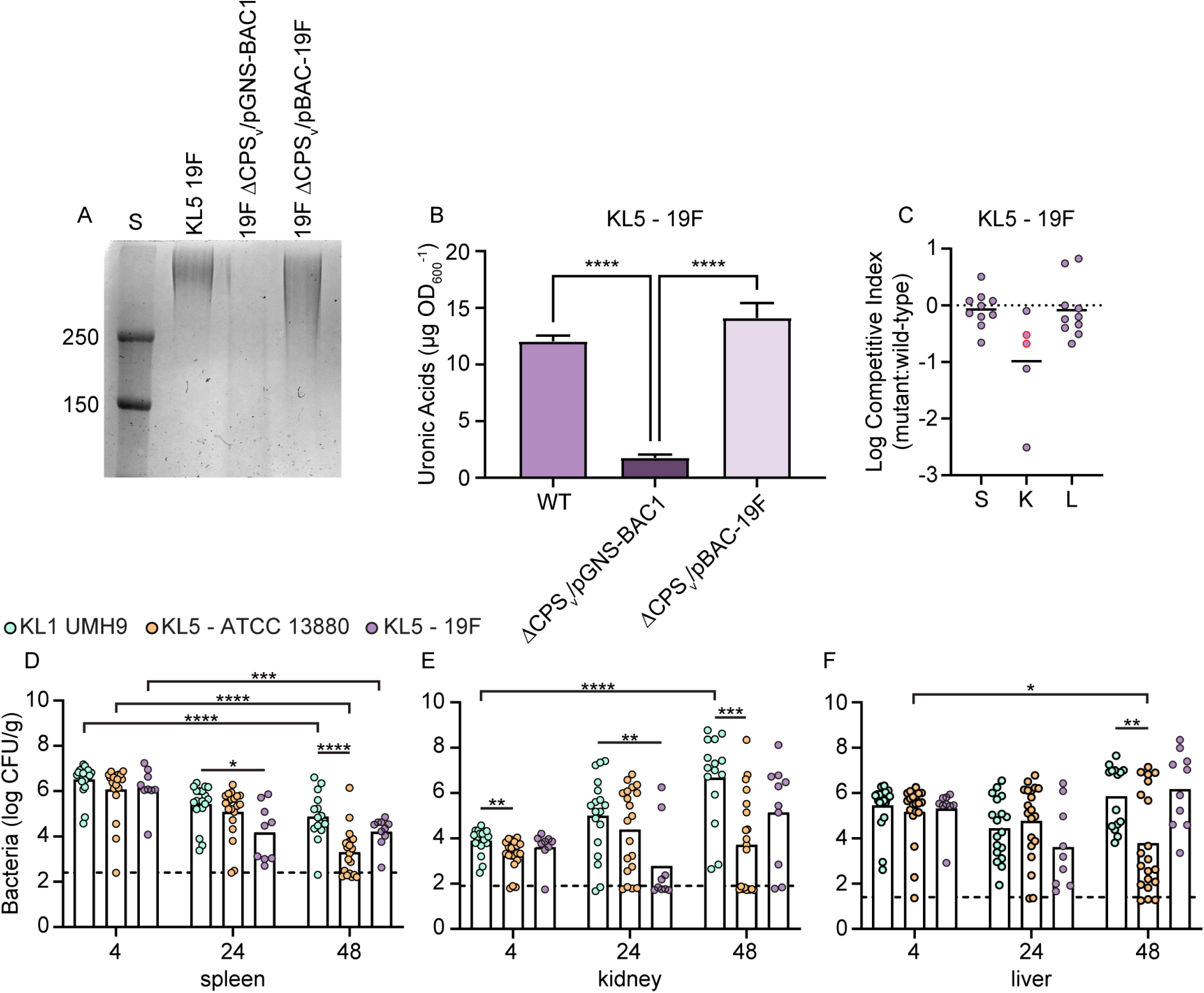
CPS does not contribute to fitness of KL5 strain 19F. A. Total polysaccharides prepared from wild-type and capsule-null derivatives of KL5 strain 19F were separated by SDS-PAGE and stained with alcian blue. The 19F capsule mutant harbored the pGNS-BAC1 vector control plasmid or recombinant plasmid containing a cloned copy of 19F KL. Pre-stained protein molecular weight standards (S) of known molecular weight are shown (kDa). B. Cell-associated uronic acids were quantitated from 19F strains in comparison to a glucuronic acid standard curve. Statistical significance was calculated by one-way ANOVA with Dunnett’s multiple comparisons test relative to 19F ΔCPS_v_/pGNS-BAC1. C. Strains 19F and 19F ΔCPS_v_ were inoculated at an equal ratio into C57BL/6J mice (n=10) and CI was determined from bacteria recovered from spleen (S), kidney (K), and liver (L). Red outlined symbols indicate CI for which mutant bacteria were recovered at or below the limit of detection. Points below the dotted line represent samples in which the mutant strain was outcompeted by wild-type bacteria. Mean log competitive indices (solid lines) were not significantly different from the hypothesized value of zero representing neutral fitness, as determined by one-sample t-test. D-F. Mice from at least two independent cohorts (n=10-20) were inoculated with the indicated strains via the TVI route and bacterial burdens in the spleen, kidneys, and liver were determined. The mean of log transformed numbers of viable bacteria recovered are indicated by the bars and dashed lines denote the highest value among samples that were at or below the limit of detection. Statistical significance was assessed by one-way ANOVA with Dunnett’s multiple comparisons test: *, Adj. P < 0.05; **, Adj. P < 0.01; ***, Adj. P < 0.001; ****, Adj. P < 0.0001.

### Sialylated CPS protect *S. marcescens* during macrophage interactions

We previously concluded that KL1 CPS has anti-phagocytic properties based on data demonstrating that a KL1 acapsular derivative was internalized more readily by the U937 monocytic cell line compared to wild-type bacteria (17). To determine whether CPS from other KL types also inhibited cellular uptake, intracellular bacteria were quantitated after incubation with murine bone marrow-derived macrophages (BMDM). The relative number of viable intracellular acapsular mutants was compared to wild-type bacteria at three time points following treatment with gentamicin to kill extracellular bacteria and calculated as an internalization index (Fig. 6A). For both KL1 and KL2 sialylated capsule types, higher numbers of viable and internalized acapsular bacteria were recovered compared to wild-type (Fig. 6B and C). Therefore, both KL1 and KL2 CPS contribute to macrophage phagocytosis resistance. In contrast, none of the non-sialylated CPS KL types exhibited a significant difference under the same conditions, as evidenced by neutral internalization indices for KL3, KL4, and KL5 bacteria (Fig. 6D-F). The comparative lack of CPS-dependent phagocytosis resistance for the KL3 and KL4 clinical strains, in particular, suggests an important role for sialylated *S. marcescens* CPS in innate immune interactions and may be one of multiple contributing factors to the successful adaptation of these lineages to infection.

**Figure 6.**
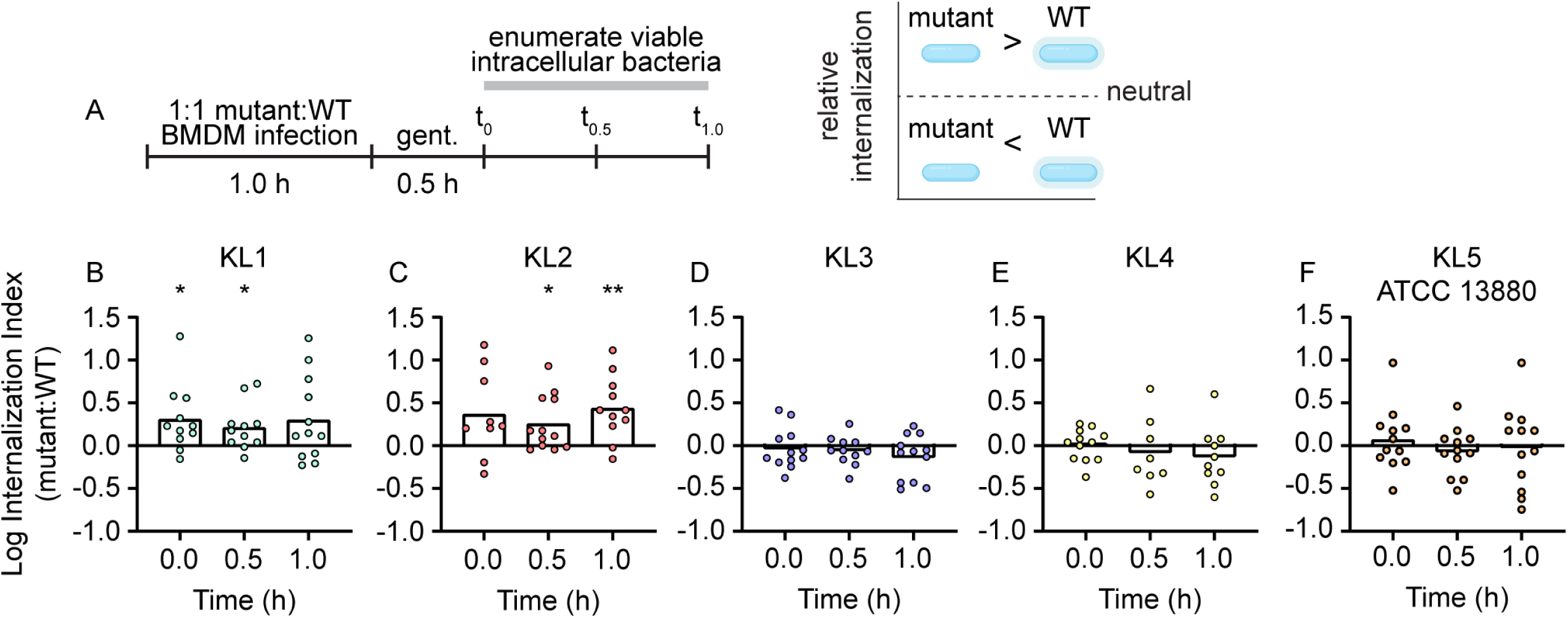
Sialylated CPS protects *S. marcescens* from macrophage internalization. A. Schematic for design and interpretation of BMDM competition infection experiments. B-E. Murine BMDM were co-infected at a 1:1 ratio with wild-type and capsule mutant derivatives of each clade followed by enumeration of viable intracellular bacteria. The relative number of intracellular mutant and wild-type (WT) bacteria was calculated as an internalization index. Bars represent the mean of log transformed internalization indices and significant deviation from the hypothetical value of zero representing equivalent internalization was determined by one-sample t-test: *, P < 0.05; **, P < 0.01.

### KL cloning and non-native CPS synthesis

KL1-4 and two KL5 loci were cloned into the bacterial artificial chromosome (BAC) pGNS-BAC1 (22). The cloned regions ranged from 15-23 kb in length (Table 1) and consisted of the entire intergenic sequence upstream of *galU*, the five-gene conserved KL region, and all clade-specific KL open reading frames (Fig. S1A). Transformation of the pBAC-KL plasmids into their respective capsule mutant strains resulted in complete restoration of extracellular uronic acid production for strains KL1-4 (Fig. 7A-D) and the 19F strain of KL5 (Fig. 5B), but not ATCC 13880 (Fig. 7E). Thus, the engineered pBAC-KL constructs are functional and sufficient to restore CPS production in their cognate strains.

**Figure 7.**
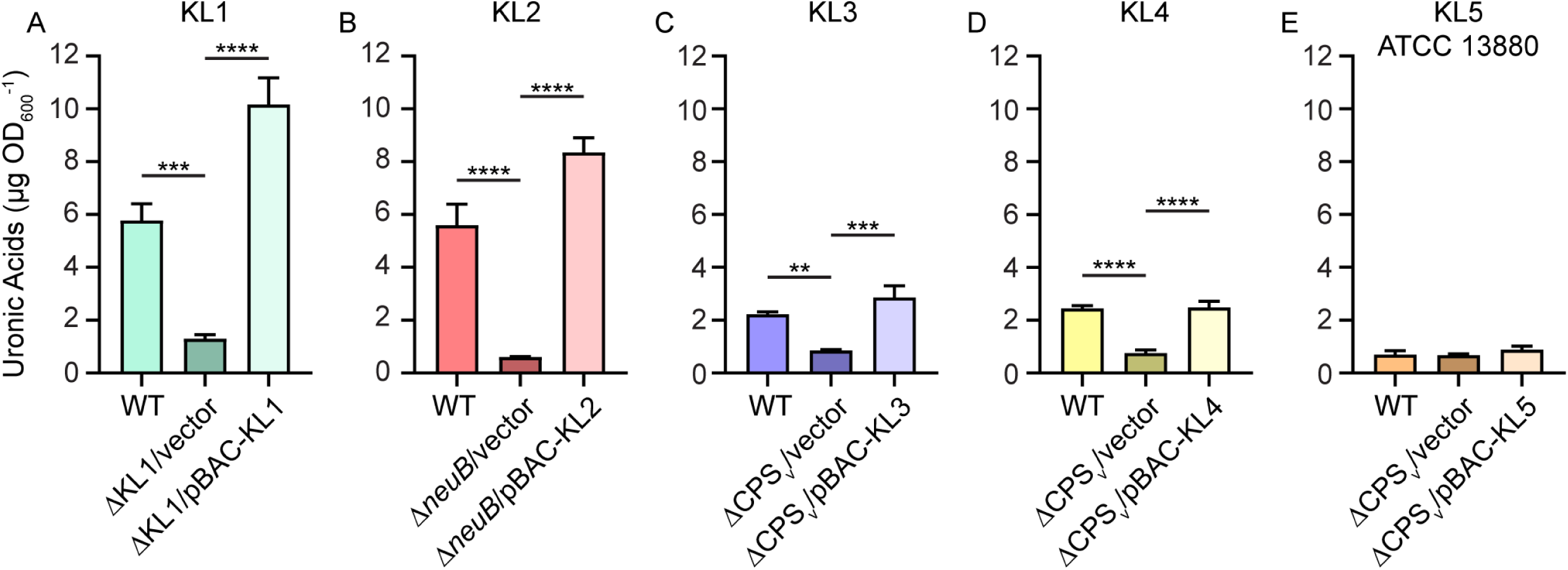
Genetic complementation of KL mutations. A-E. Capsule production by wild-type (WT) and capsule mutant *S. marcescens* strains representative of clades KL1-5 was measured by quantitating extracellular uronic acids and based on a standard curve of glucuronic acid. Statistical significance was assessed relative to mutant strains harboring the vector control plasmid by one-way ANOVA with Dunnett’s multiple comparisons test: **, Adj. P < 0.01; ***, Adj. P < 0.001; ****, Adj. P < 0.0001.

Heterologous CPS synthesis was first attempted in KL1 UMH9 because it is the strain that we have characterized most extensively. A second acapsular KL1 derivative was generated for this purpose that harbored a deletion of the entire KL1 from the five-gene conserved region (*galU*, *galF, wza*, *wzb*, *wzc*) through the clade-specific variable locus (Fig. S1A). BAC constructs harboring KL1-4 were transformed into the UMH9 ΔKL1 strain and CPS production was quantitated via uronic acids. Unexpectedly, none of the BAC constructs harboring KL2-4 yielded significant increases in uronic acids compared to the ΔKL1 vector control (Fig. 8A). Total extracellular polysaccharides were also isolated from these strains and resolved by SDS-PAGE. Consistent with the uronic acid quantitation, each of the pBAC-KL constructs was able to restore production of CPS when introduced into the native acapsular strains but not in the ΔKL1 background (Fig. 8B-E). We hypothesized that the Wzi protein, required for surface attachment of CPS and encoded outside the KL (17), may be involved in strain specific CPS display. However, a cloned copy of the KL3 *wzi* gene expressed in the ΔKL1/pBAC-KL3 strain does not restore production of either cell-free or cell-associated uronic acids in this background (Fig. S5A) but does restore surface association of KL1 CPS in a KL1 Δ*wzi* mutant (Fig. S5B). Therefore, the inability to heterologously produce CPS in KL1 UMH9 is due to a presently unknown limitation but may occur prior to polysaccharide surface translocation and attachment.

**Figure 8.**
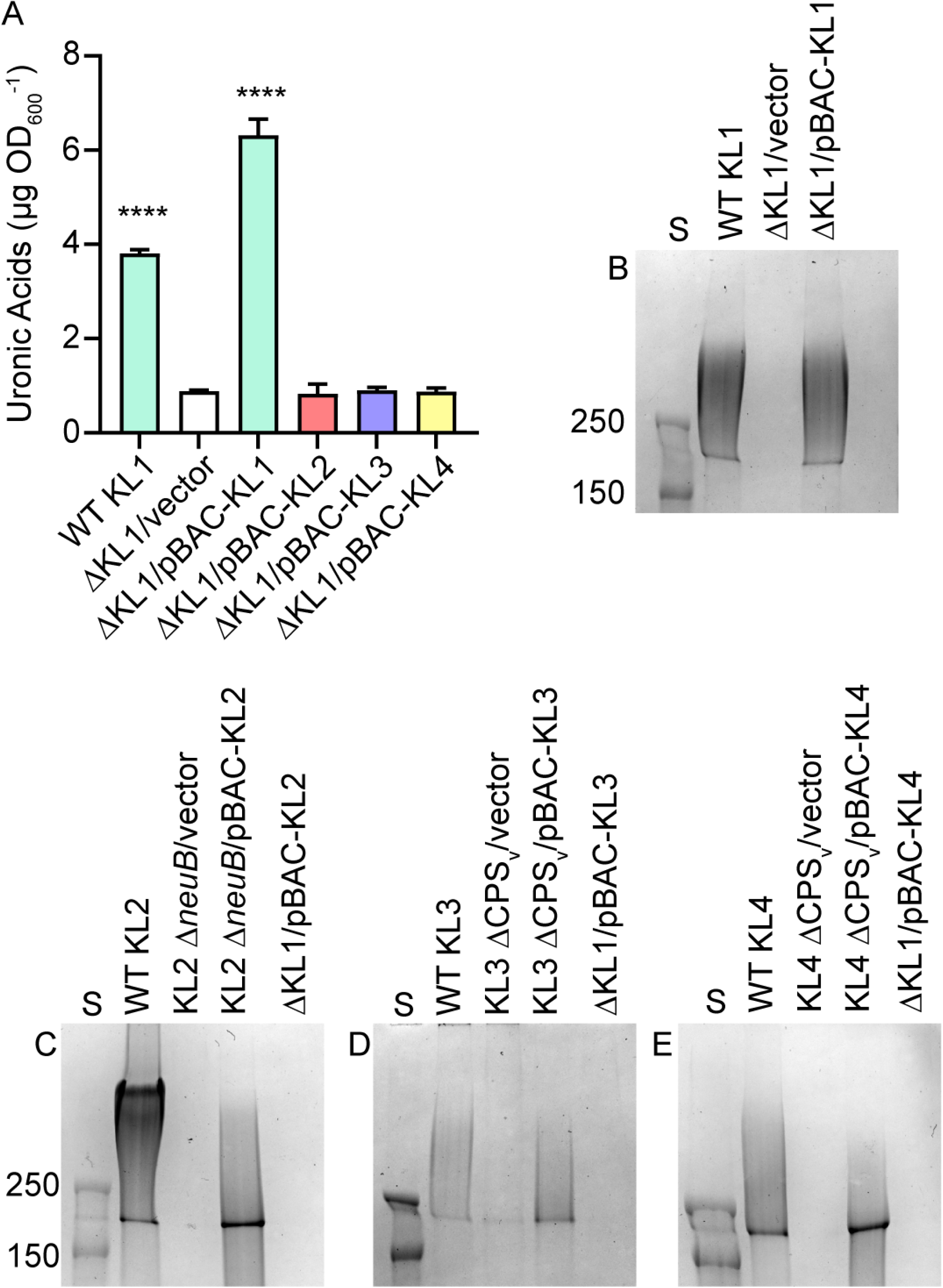
Synthesis of non-native CPS is restricted in KL1. A. Extracellular uronic acids were quantitated from the wild-type KL1 strain and a KL1 deletion mutant (ΔKL1) harboring the vector control plasmid pGNS-BAC1 or plasmids containing the KL1-4 regions. Uronic acids were quantitated in comparison to a glucuronic acid standard curve. Statistical significance was determined using ordinary one-way ANOVA with Dunnett’s multiple comparisons test against the negative control strain: ****, Adj. P<0.0001. B-E. Total bacterial polysaccharides from wild-type (WT) and capsule-null derivatives were separated by SDS-PAGE and stained with alcian blue. Capsule mutants harbored either the pGNS-BAC1 vector control plasmid or a recombinant plasmid with a cloned copy of the native KL. Recombinant KL plasmids were also expressed from a capsule-null mutant derivative of KL1 (ΔKL1). Pre-stained protein molecular weight standards (S) were electrophoresed on each gel with molecular weights shown in kDa.

To determine if non-native CPS production was possible in other lineages, BAC constructs containing KL1 and KL2 were transformed into the KL3 and KL4 ΔCPS_v_ mutants. KL1 and KL2 CPS were quantitated at or above the level of wild-type strains and the native pBAC-KL3 or pBAC-KL4 complemented ΔCPS_v_ mutants as measured by total cell-associated plus extracellular uronic acids in both backgrounds (Fig. 9A and 9C). However, only a minor fraction of KL1 CPS was surface associated compared to CPS from the other BAC constructs (Fig. 9B and 9D). Since abundant non-native KL1 CPS is likely released from the surface in these scenarios, further KL1 genetic combinations were not pursued. The pBAC-KL2 construct in contrast yielded surface-associated KL2 CPS at levels similar to the native pBAC-KL3 and pBAC-KL4 constructs. Sialic acids were next quantitated as an additional measure of KL2 CPS in KL3 and KL4 strains. Wild-type KL3 and KL4 strains yielded background levels of sialic acids by the thiobarbituric acid assay and both strains carrying pBAC-KL2 yielded a significant increase in extracellular sialic acids compared capsule mutant bacteria harboring the vector control plasmid (Fig. 9E). These combined results demonstrate that sialylated and surface-associated KL2 CPS can be synthesized in both KL3 and KL4 strains.

**Figure 9.**
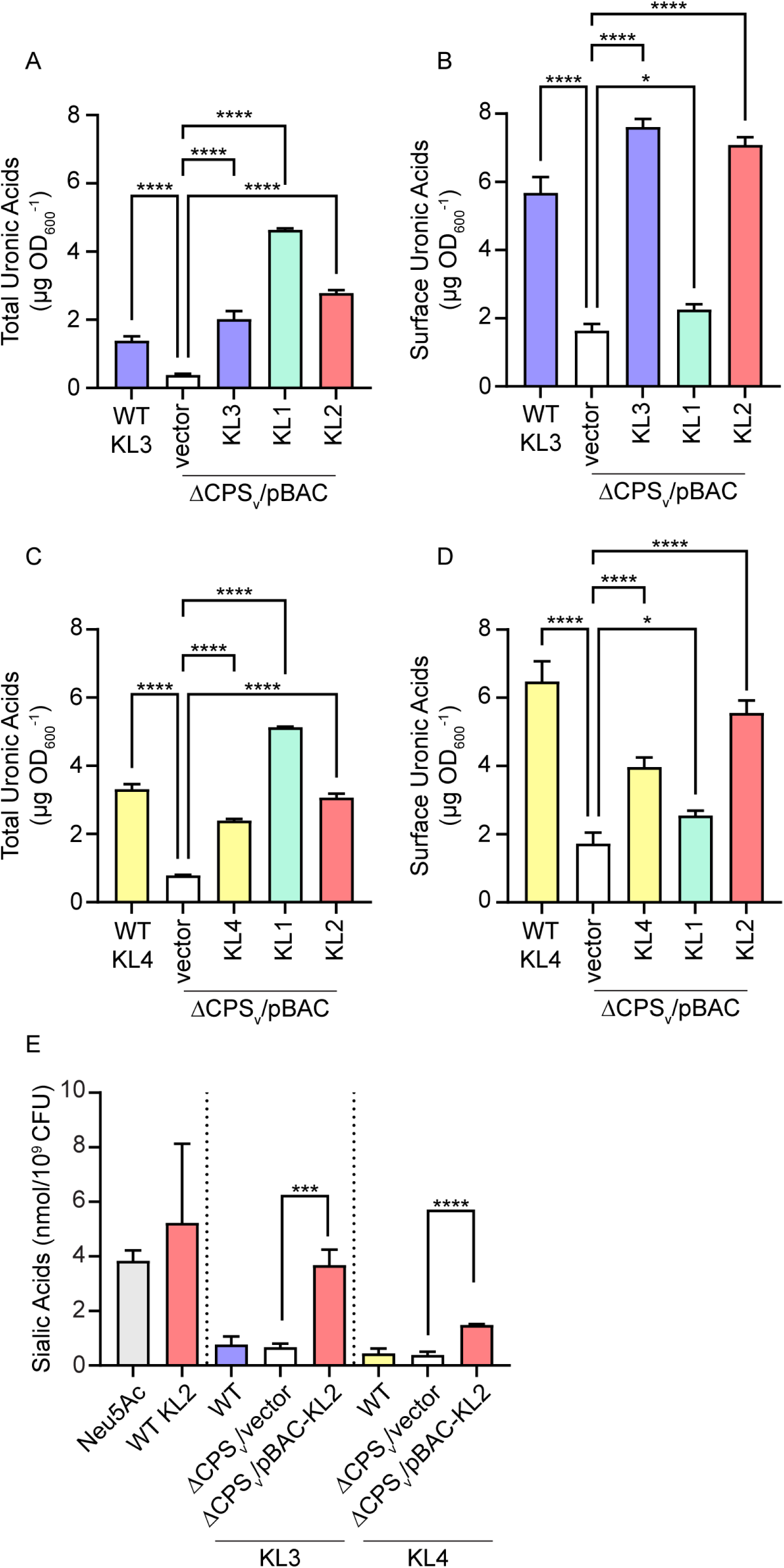
Non-native synthesis of sialylated CPS in KL3 and KL4 strains. A-D. Total uronic acids (A and C) and surface-associated uronic acids (B and D) were quantitated from wild-type (WT) and ΔCPS_v_ mutant strains. Mutant bacteria harbored either the control plasmid pGNS-BAC (vector) or recombinant BAC plasmids containing the indicated KL. Quantitation was based on a standard curve with glucuronic acid. Statistical significance was assessed by one-way ANOVA with Dunnett’s multiple comparisons test against ΔCPS_v_ mutants carrying the vector control plasmid. Adj. P: *, <0.05; **, <0.01; ***, <0.001; ****, <0.0001. E. Sialic acids were quantitated by thiobarbituric acid assay and normalized by CFU between strains. Purified Neu5Ac and CPS from wild-type KL2 bacteria were used as positive controls. KL3 and KL4 ΔCPS_v_ mutants harbored either the empty pGNS-BAC plasmid or recombinant plasmids expressing KL2. Statistical analysis was constrained within the dotted lines and was performed as described for panels A-D.

### Non-native KL2 CPS limits macrophage internalization

To further investigate the specific impact of *S. marcescens* sialylated CPS on macrophage interactions, immunofluorescence microscopy was used to quantitate intracellular and extracellular bacteria associated with BMDM (Fig. 10A). KL1 bacteria with and without the native KL1 CPS were first analyzed to establish the approach. Both wild-type and the ΔKL1 mutant had a similar number of total bacteria (extracellular + intracellular) associated with BMDM on a per-cell basis (Fig. 10B). However, a significantly higher proportion of these bacteria were extracellular for wild-type whereas the ΔKL1 derivative was predominantly intracellular (Fig. 10C and D, Fig. S6). These results confirm the previous gentamicin protection assay findings (Fig. 6) and solidify the role of KL1 CPS in resisting macrophage phagocytosis. Genetic complementation with pBAC-KL1 restored the extracellular-to-intracellular relationship observed for the wild-type strain (Fig. 10E), but substantially reduced the total number of bacteria per BMDM (Fig. 10B), potentially due to CPS hyperproduction (Fig. 7A).

**Figure 10.**
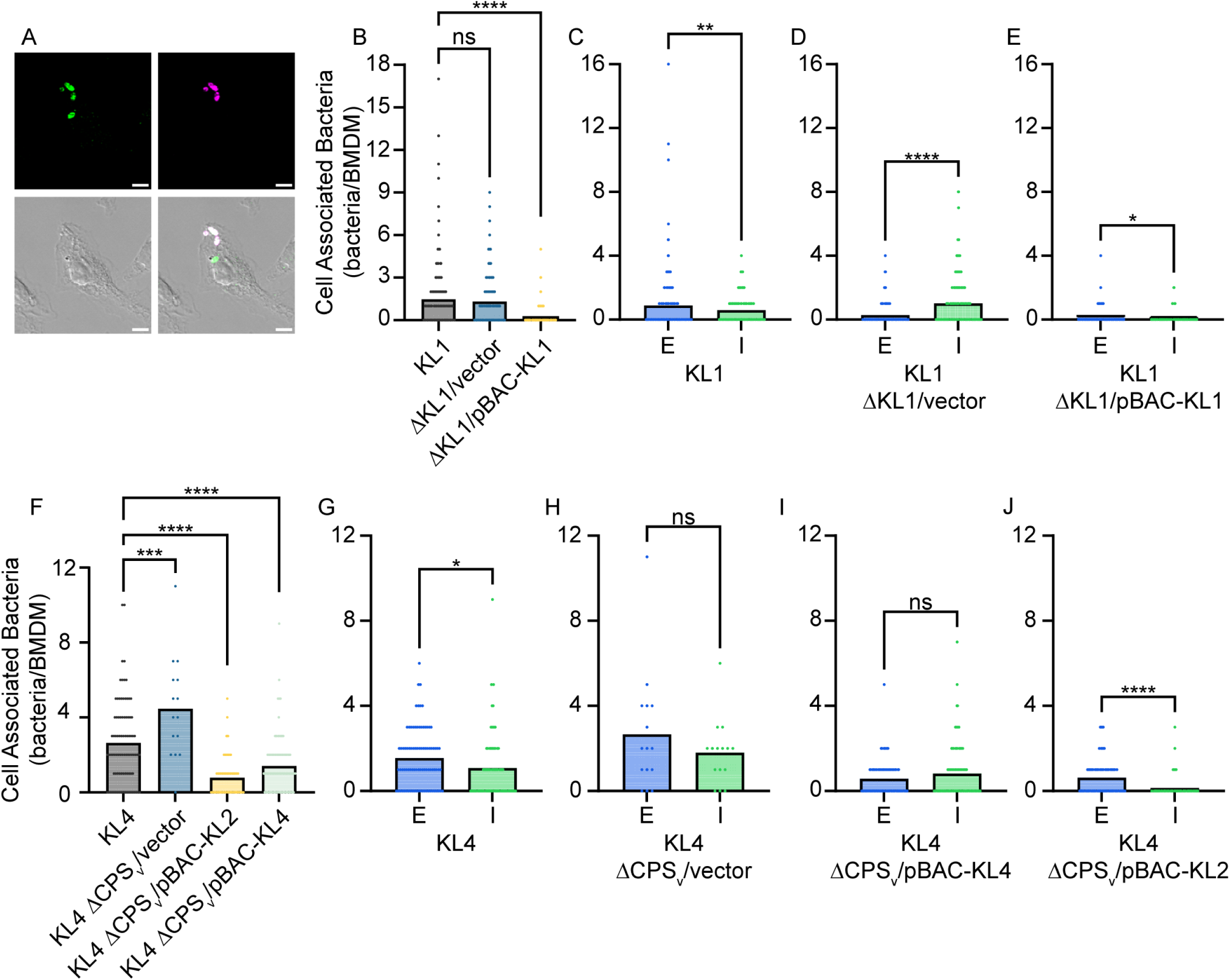
Ectopic expression of KL2 genes confers phagocytosis resistance to a non-sialylated strain. A. Representative images of BMDM infected with the wild-type KL1 strain for 60 min followed by differential immunofluorescence and microscopy. Extracellular bacteria were labeled with an AlexaFluor-647 conjugated secondary antibody, BMDM were then permeabilized and all bacteria were exposed to an AlexaFluor-488 conjugated secondary antibody. Extracellular bacteria fluoresce in both channels and appear white in the composite image while intracellular bacteria appear green. Scale bars are 5 µm. B and F. Quantitation of total BMDM-associated bacteria (extracellular + intracellular). Points represent the number of bacteria associated with individual BMDM cells and statistical significance was assessed by one-way ANOVA with Dunnett’s multiple comparisons test against the KL1 and KL4 wild-type strains, respectively. Adj. P values: ***, <0.001; ****, <0.0001. C-E and G-J. Quantitation of extracellular (E) and intracellular (I) BMDM-associated bacteria for KL1 and KL4 wild-type and CPS mutant derivative strains. Statistical significance was assessed by unpaired t-test. P values: *, <0.05; **, <0.01; ***, <0.001; ****, <0.0001; ns, non-significant.

The ability to synthesize KL2 CPS in both the KL3 and KL4 strains allowed us to test the hypothesis that sialylated KL2 CPS confers phagocytosis resistance to these normally non-sialylated lineages. Loss of the native KL3 and KL4 CPS significantly increased the per-cell number of total BMDM-associated bacteria compared to the parental strains (Fig. 10F and S7) but as expected, the relationship between the number of extracellular and intracellular bacteria remained largely unchanged between wild-type and ΔCPS_v_ mutant bacteria (Fig. 10G and H, Fig. S7B and C). Thus, the KL3 and KL4 CPS have minimal impact on BMDM phagocytosis, as demonstrated in Fig. 6. Unfortunately, complementation of the KL3 ΔCPS_v_ mutant with either pBAC-KL3 or pBAC-KL2 significantly reduced the overall detectable number of cell-associated bacteria to a level that prevented reliable assessment of intracellular and extracellular trends (Fig. S7). This was not the case for KL4 and in fact, complementation with pBAC-KL4 confirmed that native KL4 synthesis does not result in significant differences between the number of intracellular and extracellular bacteria per cell (Fig. 10I), similar to the KL4 ΔCPS_v_ mutant harboring the vector control (Fig. 10H). In contrast, KL4 ΔCPS_v_ harboring pBAC-KL2 shifted localization to a significant majority of extracellular bacteria compared to nearly undetectable numbers of intracellular bacteria (Fig. 10J). Sialylated KL2 CPS is therefore capable of restricting macrophage phagocytosis of both its native strain and a normally non-sialylated *S. marcescens* lineage.

## DISCUSSION

In this study, *S. marcescens* strains isolated from clinical and non-clinical sources were assayed for survival characteristics during infection. These strains were selected as representatives of five sequence-defined capsule clades. While all strains demonstrated an ability to infect mice using two bacteremia models, significant differences in organ colonization and immune cell interactions were observed. Though we attempted to capture a range of diverse isolates within the species, one acknowledged limitation of the study is the use of a single KL representative in most cases. Nonetheless, capsule is established here as a critical fitness determinant for each of the clinical clades assessed. Given the mounting genomic evidence distinguishing *S. marcescens* clinical lineages from environmental isolates (12, 14, 15), it’s likely that other factors also contribute the infection fitness of healthcare-associated strains and our ongoing work aims to identify and characterize such factors encoded within these clinical accessory genomes. Strains belonging to the environmental capsule clade KL5 generally exhibited a lower infection capacity and less dependence on CPS in our models. For type strain ATCC 13880, these observations could be attributed to an inability to synthesize CPS. Repeated laboratory passage of encapsulated bacteria can result in spontaneous loss of capsule due to mutation, but whether this is the case for ATCC 13880 remains to be determined. However, KL5 strain 19F yields abundant CPS yet exhibited no significant fitness cost when capsule was lost, supporting the conclusion that KL5 CPS plays a minor role bacteremia.

Two approaches were used to demonstrate that sialic acid containing CPS of KL1 and KL2 *S. marcescens* had the greatest protective effect against BMDM internalization. We hypothesize that KDN and/or Neu5Ac uniquely associated with these capsule clades (17) may therefore have a specific role in influencing *Serratia*-macrophage interactions. While such molecular interactions, particularly for the more abundant and understudied KDN component, have yet to be demonstrated for *S. marcescens*, this hypothesis is indirectly supported by experimentally established roles for Neu5Ac-mediated modulation of innate immune cells in other systems (23–25). It’s notable that non-sialylated clinical CPS types also have a meaningful, but perhaps different, role in infection as demonstrated by the significant fitness cost of acapsular KL3 and KL4 derivatives. With exception of the aforementioned *wzi* gene, each *S. marcescens* KL was expected to be sufficient for type-specific CPS synthesis; however, the limited ability or inability to produce non-native CPS in some strains indicates that additional unknown factors are required. KL1 bacteria in particular failed to synthesize any of the other CPS, including the closely related CPS of KL2, and KL1 CPS was only poorly surface-associated in the other lineages.

In our previous KL comparison, the KL1 and KL2 lineages had the greatest number of representatives in our genome cohort and both were overwhelmingly comprised of infection isolates (17). In the context of the comprehensive *S. marcescens* genomic architecture published by Ono *et al*. (12), the UMH9 KL1 and gn773 KL2 strains characterized here both segregate into clade 1. This is notable because clade 1 *S. marcescens* are almost exclusively hospital associated or clinical isolates, have a higher number of antimicrobial resistance alleles, and encode a distinct set of accessory genes compared to other lineages. Furthermore, of the 215 strains identified as either KL1 or KL2 in our study, 188 were also included in the Ono study and all of them were assigned to the clade 1 genomic lineage. This observation independently confirms our conclusion that KL1 and KL2 CPS are a differential component of these infection-adapted *S. marcescens* and together with the results reported here indicate that sialylated CPS contribute to the niche-specific characteristics that provide these isolates with a selective advantage during infection.

## MATERIALS AND METHODS

### Bacterial strains and culture conditions

The *S. marcescens* strains used in this study are listed in Table 1. *Escherichia coli* DH10B and DH5α were routinely used for cloning purposes. DH5α harboring helper plasmid pRK2013 (26) or *E. coli* BW29427 (B. Wanner, unpublished) cultured in 0.3 mM diaminopimelic acid were used as donor strains for conjugation. Bacteria were cultured in either LB medium (27) with or without 20 mM glucose or M9 (28) medium supplemented with 1 mM MgSO_4_, 36 µM FeSO_4_, 100 µM CaCl_2_ and 20 mM glucose. Antibiotics for bacterial culture were used at the following concentrations: kanamycin, 50 µg/mL; hygromycin, 200 µg/mL; spectinomycin, 100 µg/mL; gentamicin, 10 and 20 µg/mL; and ampicillin, 100 µg/mL.

### Generation of mutants

The *S. marcescens* ATCC 13880 and UMH7 KL mutations (Table 1) were constructed by recombineering as previously described (18, 29). Briefly, the *nptII* gene from pKD4 (30) was PCR-amplified with oligonucleotides possessing ∼50-bp of 5′ sequence homology to the targeted CPS_v_ region. Electrocompetent recipient strains harboring pSIM18 or pSIM19 (31) were transformed with DpnI-treated PCR products. Kanamycin-resistant transformants were genotyped by PCR and sequenced, then cured of pSIM plasmids prior to use in phenotypic assays. KL mutations in strains UMH11, gn773, and 19F (Table 1) were accomplished via allelic exchange with pTOX11 (32) derivatives as previously described (17). Plasmids were constructed from PCR-amplified fragments using NEB HiFi Assembly and harbored ∼700 bp of homologous sequence flanking the *nptII* allele to facilitate recombination. The unmarked ΔKL1 mutation in strain UMH9 was generated using a similar approach, but did not include the *nptII* insertion. Allelic exchange was performed following conjugation of pTOX plasmids from *E. coli* donor strain BW29427 into recipient *S. marcescens* strains as previously described (17, 32). The presence of mutant alleles was confirmed by PCR amplification and sequencing. Transconjugants were also assessed by PCR to ensure Mu phage was not transferred from the BW29427 donor (33). All primer sequences used for PCR amplification in the procedures described above are listed in Table S1.

### Genetic complementation

KL sequences were cloned into BAC vector pGNS-BAC1 (22). Each KL region with upstream intergenic sequence was PCR amplified in two fragments using the primers listed in Table S1. Fragments were then cloned into HindIII-digested pGNS-BAC1 using HiFi DNA Assembly and transformed into electrocompetent *E. coli* DH10B. Recombinant pGNS-BAC1 plasmids were purified by alkaline lysis and whole-plasmid sequences were determined by Nanopore (SNPsaurus). The resulting pBAC-KL plasmids (Table 1) were transferred to *S. marcescens* recipient strains via tri-parental mating using helper plasmid pRK2013 (26). Ampicillin and gentamicin were used to select for the loss of *E. coli* donor strains and the presence of pBAC-KL plasmids in *S. marcescens*, respectively.

### Quantitation of uronic acids and polysaccharide analysis

Extracellular uronic acids of *S. marcescens* were measured using previously described methods (17, 34, 35). Measurements were based on a standard curve of glucuronic acid and normalized to culture optical density (600 nm). Results are the means from three biological replicates and are representative of at least two independent experiments. Isolation of *S. marcescens* extracellular polysaccharides and visualization by SDS-PAGE was also performed according to published methods (17, 36) and the results are representative of at least two independent experiments.

### Human serum exposure

Bacterial viability following a 90-minute incubation in 40% pooled human serum (Innovative Research) was determined as previously described (17). Results are the means from three biological replicates and are representative of three independent experiments.

For the attempted selection of ATCC 13880 CPS synthesis, wild-type and the ΔCPS_v_ derivative were passaged every 24 h in LB medium or LB medium supplemented with increasing concentrations of pooled normal human serum (5%, 10%, 20%) over the course of 72 h. The presence of CPS was assessed via negative stain with Maneval’s reagent (37) and visualized using a Nikon Ti2 widefield microscope with a 100x objective (University of Michigan Microscopy Core). KL2 bacteria harvested from LB agar was used as a positive control.

### Quantitation of sialic acids

Sialic acids were quantitated from *S. marcescens* strains by thiobarbituric assay as previously described (17, 38). Extracellular polysaccharides were subjected to acid hydrolysis with 0.1 N HCl at 80°C for 60 min and bacteria were subsequently removed by centrifugation. Hydrolyzed solutions were subjected to periodate oxidation according to the protocol and reacted with thiobarbituric acid. Chromophore extraction was performed with an equal volume of cyclohexanone and absorbance was measured at 549 nm. The amount of sialic acid was determined with the following formula: (Volume (mL) prior to extraction x OD_549_) / 57 = μmole Neu5Ac equivalents. The amount of sialic acid detected was normalized to the bacterial culture density and 100 μM Neu5Ac served as the positive control. Results are reported as the mean from three biological replicates and representative of two independent experiments.

### Murine infections

Murine infections were performed with protocols approved by the University of Michigan Institutional Animal Care and Use Committee and in accordance with Office of Laboratory Animal Welfare guidelines. For TVI mono-infections, male and female 7-8 week-old C57BL/6J mice (Jackson Laboratories) were inoculated with 0.1 mL bacterial suspensions in PBS containing a target dose of 5×10^6^ CFU. For TVI competition infections, wild-type bacteria were mixed at a 1:1 ratio with mutant strains and delivered at a dose of 5×10^6^ total CFU. Mice were euthanized 24 h post-infection, unless otherwise specified, and the spleen, liver, and kidneys were harvested and homogenized. Bacterial counts of the inoculum (input) and organ homogenates (output) were determined by plating serial dilutions on LB agar with or without kanamycin. The CI was determined by the following calculation: (CFU_mutant_/CFU_wildtype_)^output^/(CFU_mutant_/CFU_wildtype_)^input^. A log transformed CI of less than zero indicates a competitive disadvantage for mutant bacteria compared to the parental strain.

For the bacteremic pneumonia model, 0.05 ml bacterial suspensions were delivered to the retropharyngeal space of anesthetized 7-8 week-old mice at a target dose of 1×10^7^ total CFU. Mice were euthanized 24 h post-inoculation and the spleen, liver, kidneys, and lung were harvested. CI was determined as described above.

### Gentamicin protection assays

Isolation and propagation of BMDM was accomplished using established protocols (39). Monocytes from the femur and tibia bone marrow of 7- to 8-week-old C57BL/6J mice were diluted to 1×10^6^ cells/mL in medium containing 15% L929 cell supernatant. At 7 days post-harvest, BMDM were dissociated from wells with ice-cold 2 mM EDTA in DPBS and collected by centrifugation. BMDM were seeded into 96-well flat bottom plates at 1×10^5^ cells/well and incubated at 37°C at 5% CO_2_ for 24 h prior to inoculation with bacteria. Wild-type and kanamycin-resistant mutant strains were mixed in 1:1 ratio and added to BMDM at target MOI of 20. Plates were centrifuged briefly and incubated at 37°C for 60 min in 5% CO_2_. The medium was then aspirated and wells were washed with DPBS. DMEM containing 10% FBS and 100 µg/mL gentamicin was added to wells for 30 min at 37°C in 5% CO_2_ followed by removal of gentamicin-containing medium and washing. For time point zero, BMDM were exposed to 1% saponin at 37°C for 10 min, mixed with 0.1 mL LB medium, then serially diluted and plated on LB agar plate with and without kanamycin for CFU determination. All other time points were incubated in medium containing 10 µg/mL gentamicin until permeabilization and CFU determination. Internalization indices from two independent experiments were calculated as described for CI with internalized bacteria substituting for the infection output parameter.

### Immunofluorescence

BMDM were allowed to adhere to glass coverslips overnight and then infected as described for the gentamicin protection assays with single *S. marcescens* strains. After 60 min incubation, coverslips were washed with PBS and fixed with 4% paraformaldehyde. PBS with 10% goat serum was used to block coverslips prior to incubation with a 1:100 dilution of the primary antibody, an anti-*E. coli* polyclonal antibody (Invitrogen, AB_780488) that cross-reacts with *S. marcescens*. After washing, a 1:400 dilution of goat anti-rabbit secondary antibody conjugated to Alexa Fluor 647 (Invitrogen, AB_2535813) was applied. Coverslips were thoroughly washed and fixed again in paraformaldehyde, followed by permeabilization of BMDM with 0.2% saponin and re-application of the primary antibody. A 1:400 goat anti-rabbit secondary antibody conjugated to Alexa Fluor 488 (Invitrogen, AB_143165) was applied to differentiate internalized bacteria. Imaging was conducted on a Nikon N-SIM A1R confocal microscope (University of Michigan Microcopy Core) using a 60x objective. All image analysis was performed in the Fiji distribution of ImageJ (40). Intracellular and extracellular bacteria were differentiated by single (488 nm) or dual fluorescence, respectively, and only cell-associated bacteria were counted. Each strain was assayed in at least two independent experiments and images were collected from a minimum of 15 fields per coverslip.

## ACKNOWLEDGEMENTS

The authors would like to thank Mark Liles for kindly sharing the pGNS-BAC1, Nancy Moran for providing *S. marcescens* strains, and Caity Holmes for providing L929 cells and guidance regarding BMDM propagation. This work was supported by Public Health Service award AI148767 to M. T. A. from the National Institutes of Health and AI134731 from the National Institutes of Health to H.L.T.M. and M.A.B. The funders had no role in study design, data collection and analysis, or preparation of the manuscript.

## REFERENCES

1. Wisplinghoff H, Bischoff T, Tallent SM, Seifert H, Wenzel RP, Edmond MB. 2004. Nosocomial bloodstream infections in US hospitals: analysis of 24,179 cases from a prospective nationwide surveillance study. Clin Infect Dis 39:309–17.

2. Sader HS, Streit JM, Carvalhaes CG, Huband MD, Shortridge D, Mendes RE, Castanheira M. 2021. Frequency of occurrence and antimicrobial susceptibility of bacteria isolated from respiratory samples of patients hospitalized with pneumonia in Western Europe, Eastern Europe and the USA: results from the SENTRY Antimicrobial Surveillance Program (2016-19). JAC Antimicrob Resist 3:dlab117.

3. Antimicrobial Resistance Collaborators. 2022. Global burden of bacterial antimicrobial resistance in 2019: a systematic analysis. Lancet 399:629–655.

4. GBD Antimicrobial Resistance Collaborators. 2022. Global mortality associated with 33 bacterial pathogens in 2019: a systematic analysis for the Global Burden of Disease Study 2019. Lancet 400:2221–2248.

5. Raymond J, Aujard Y. 2000. Nosocomial infections in pediatric patients: a European, multicenter prospective study. European Study Group. Infect Control Hosp Epidemiol 21:260–3.

6. Dessi A, Puddu M, Testa M, Marcialis MA, Pintus MC, Fanos V. 2009. *Serratia marcescens* infections and outbreaks in neonatal intensive care units. J Chemother 21:493–9.

7. Johnson A, Watson D, Dreyfus J, Heaton P, Lampland A, Spaulding AB. 2020. Epidemiology of *Serratia* bloodstream infections among hospitalized children in the United States, 2009-2016. Pediatr Infect Dis J 39:e71–e73.

8. Mahlen SD. 2011. *Serratia* infections: from military experiments to current practice. Clin Microbiol Rev 24:755–91.

9. Raymann K, Coon KL, Shaffer Z, Salisbury S, Moran NA. 2018. Pathogenicity of *Serratia marcescens* strains in honey bees. mBio 9:e01649–18.

10. Patterson KL, Porter JW, Ritchie KB, Polson SW, Mueller E, Peters EC, Santavy DL, Smith GW. 2002. The etiology of white pox, a lethal disease of the Caribbean elkhorn coral, *Acropora palmata*. Proc Natl Acad Sci U S A 99:8725–30.

11. Mah-Sadorra JH, Najjar DM, Rapuano CJ, Laibson PR, Cohen EJ. 2005. *Serratia* corneal ulcers: a retrospective clinical study. Cornea 24:793–800.

12. Ono T, Taniguchi I, Nakamura K, Nagano DS, Nishida R, Gotoh Y, Ogura Y, Sato MP, Iguchi A, Murase K, Yoshimura D, Itoh T, Shima A, Dubois D, Oswald E, Shiose A, Gotoh N, Hayashi T. 2022. Global population structure of the *Serratia marcescens* complex and identification of hospital-adapted lineages in the complex. Microb Genom 8:000793.

13. Williams DJ, Grimont PAD, Cazares A, Grimont F, Ageron E, Pettigrew KA, Cazares D, Njamkepo E, Weill FX, Heinz E, Holden MTG, Thomson NR, Coulthurst SJ. 2022. The genus *Serratia* revisited by genomics. Nat Commun 13:5195.

14. Matteoli FP, Pedrosa-Silva F, Dutra-Silva L, Giachini AJ. 2021. The global population structure and beta-lactamase repertoire of the opportunistic pathogen *Serratia marcescens*. Genomics 113:3523–3532.

15. Abreo E, Altier N. 2019. Pangenome of *Serratia marcescens* strains from nosocomial and environmental origins reveals different populations and the links between them. Sci Rep 9:46.

16. Moradigaravand D, Boinett CJ, Martin V, Peacock SJ, Parkhill J. 2016. Recent independent emergence of multiple multidrug-resistant *Serratia marcescens* clones within the United Kingdom and Ireland. Genome Res 26:1101–9.

17. Anderson MT, Himpsl SD, Mitchell LA, Kingsley LG, Snider EP, Mobley HLT. 2022. Identification of distinct capsule types associated with *Serratia marcescens* infection isolates. PLoS Pathog 18:e1010423.

18. Anderson MT, Mitchell LA, Zhao L, Mobley HLT. 2017. Capsule production and glucose metabolism dictate fitness during *Serratia marcescens* bacteremia. mBio 8:e00740–17.

19. Anderson MT, Brown AN, Pirani A, Smith SN, Photenhauer AL, Sun Y, Snitkin ES, Bachman MA, Mobley HLT. 2021. Replication dynamics for six Gram-negative bacterial species during bloodstream infection. mBio 12:e0111421.

20. Holmes CL, Anderson MT, Mobley HLT, Bachman MA. 2021. Pathogenesis of Gram-negative bacteremia. Clin Microbiol Rev 34:e00234–20.

21. Bachman MA, Breen P, Deornellas V, Mu Q, Zhao L, Wu W, Cavalcoli JD, Mobley HL. 2015. Genome-wide identification of *Klebsiella pneumoniae* fitness genes during lung infection. mBio 6:e00775.

22. Kakirde KS, Wild J, Godiska R, Mead DA, Wiggins AG, Goodman RM, Szybalski W, Liles MR. 2011. Gram negative shuttle BAC vector for heterologous expression of metagenomic libraries. Gene 475:57–62.

23. Mandrell RE, Apicella MA. 1993. Lipo-oligosaccharides (LOS) of mucosal pathogens: molecular mimicry and host-modification of LOS. Immunobiology 187:382–402.

24. Carlin AF, Uchiyama S, Chang YC, Lewis AL, Nizet V, Varki A. 2009. Molecular mimicry of host sialylated glycans allows a bacterial pathogen to engage neutrophil Siglec-9 and dampen the innate immune response. Blood 113:3333–6.

25. Chang YC, Olson J, Beasley FC, Tung C, Zhang J, Crocker PR, Varki A, Nizet V. 2014. Group B *Streptococcus* engages an inhibitory Siglec through sialic acid mimicry to blunt innate immune and inflammatory responses in vivo. PLoS Pathog 10:e1003846.

26. Figurski DH, Helinski DR. 1979. Replication of an origin-containing derivative of plasmid RK2 dependent on a plasmid function provided in trans. Proc Natl Acad Sci U S A 76:1648–52.

27. Bertani G. 1951. Studies on lysogenesis. I. The mode of phage liberation by lysogenic Escherichia coli. J Bacteriol 62:293–300.

28. Miller JH. 1972. Experiments in Molecular Genetics. Cold Spring Harbor Laboratory, Cold Spring Harbor, N. Y.

29. Thomason LC, Sawitzke JA, Li X, Costantino N, Court DL. 2014. Recombineering: genetic engineering in bacteria using homologous recombination. Curr Protoc Mol Biol 106:1.16.1-39.

30. Datsenko KA, Wanner BL. 2000. One-step inactivation of chromosomal genes in *Escherichia coli* K-12 using PCR products. Proc Natl Acad Sci U S A 97:6640–5.

31. Datta S, Costantino N, Court DL. 2006. A set of recombineering plasmids for Gram-negative bacteria. Gene 379:109–15.

32. Lazarus JE, Warr AR, Kuehl CJ, Giorgio RT, Davis BM, Waldor MK. 2019. A new suite of allelic-exchange vectors for the scarless modification of proteobacterial genomes. Appl Environ Microbiol 85:e00990–19.

33. Ferrieres L, Hemery G, Nham T, Guerout AM, Mazel D, Beloin C, Ghigo JM. 2010. Silent mischief: bacteriophage Mu insertions contaminate products of *Escherichia coli* random mutagenesis performed using suicidal transposon delivery plasmids mobilized by broad-host-range RP4 conjugative machinery. J Bacteriol 192:6418–27.

34. Blumenkrantz N, Asboe-Hansen G. 1973. New method for quantitative determination of uronic acids. Anal Biochem 54:484–9.

35. Favre-Bonte S, Joly B, Forestier C. 1999. Consequences of reduction of *Klebsiella pneumoniae* capsule expression on interactions of this bacterium with epithelial cells. Infection and Immunity 67:554–561.

36. Tipton KA, Rather PN. 2019. Extraction and visualization of capsular polysaccharide from *Acinetobacter baumannii*. Methods Mol Biol 1946:227–231.

37. Maneval WE. 1934. Rapid staining methods. Science 80:292–4.

38. Warren L. 1959. The thiobarbituric acid assay of sialic acids. J Biol Chem 234:1971–5.

39. Weischenfeldt J, Porse B. 2008. Bone marrow-derived macrophages (BMM): isolation and applications. CSH Protoc 2008:pdb.prot5080.

40. Schindelin J, Arganda-Carreras I, Frise E, Kaynig V, Longair M, Pietzsch T, Preibisch S, Rueden C, Saalfeld S, Schmid B, Tinevez JY, White DJ, Hartenstein V, Eliceiri K, Tomancak P, Cardona A. 2012. Fiji: an open-source platform for biological-image analysis. Nat Methods 9:676–82.

