## Supplemental Figure 1 for "Infection characteristics among *Serratia marcescens* capsule lineages"

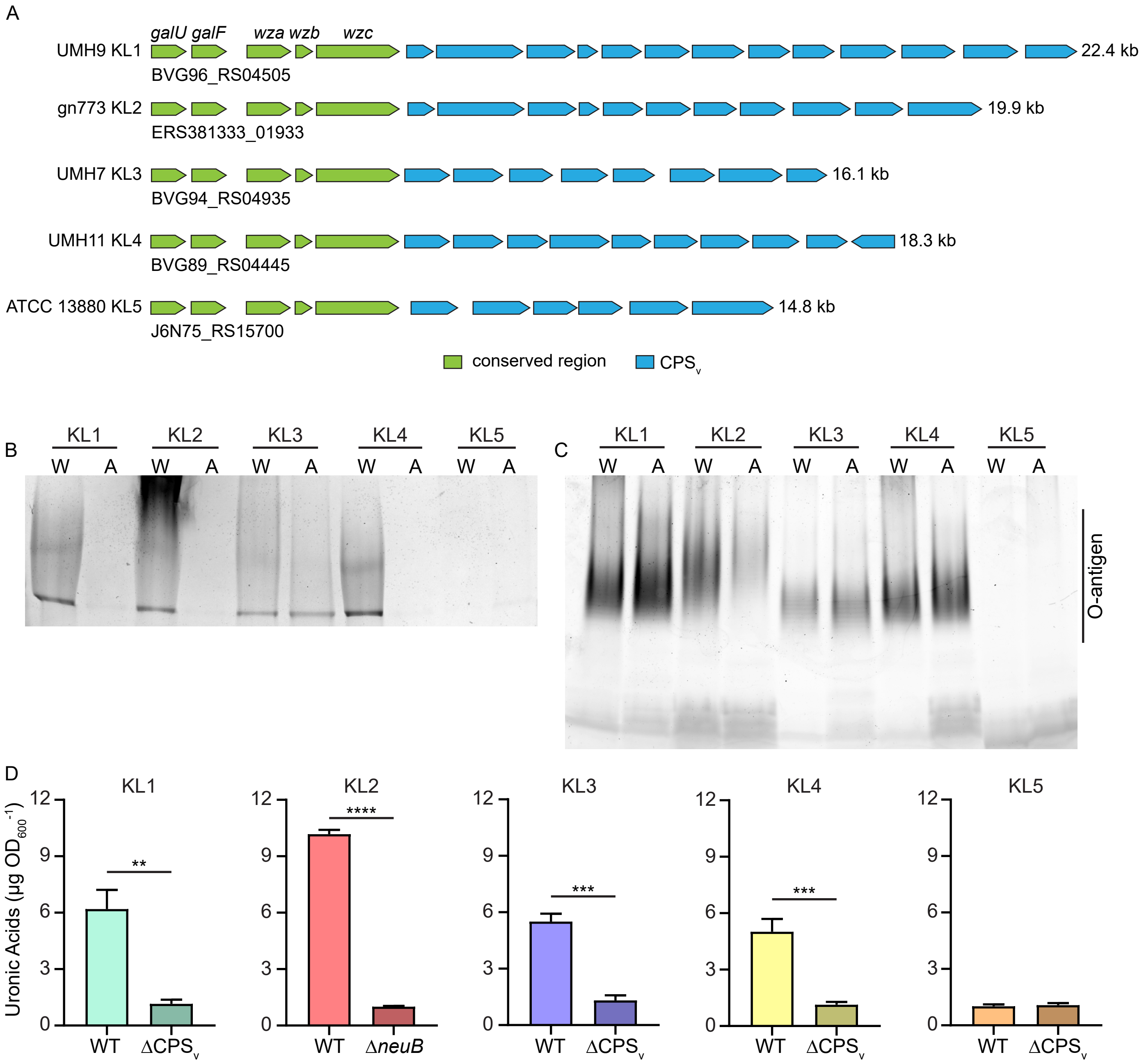


**Supplemental Figure 1. Polysaccharide phenotypes of capsule mutant strains.** A. Genetic organization of the five KL relevant to this study. The locus tag of the first open reading frame (arrows) (*galU*) in each KL is provided for reference. B. Total polysaccharides isolated from wild-type cells (W) and acapsular mutant derivatives (A) were separated by SDS-PAGE and stained with alcian blue. C. The same polysaccharides from panel A were electrophoresed with a 7-20% SDS-PAGE gel and stained with ProQ Emerald green to visualize LPS components. D. Measurement of extracellular uronic acids in wild-type (WT) and capsule mutant derivatives of KL1-5 strains. Statistical significance was assessed by unpaired t-test: **, $P < 0.01$; ***, $P < 0.001$; ****, $P < 0.0001$.
