## Supplemental Figure 2 for "Infection characteristics among *Serratia marcescens* capsule lineages"

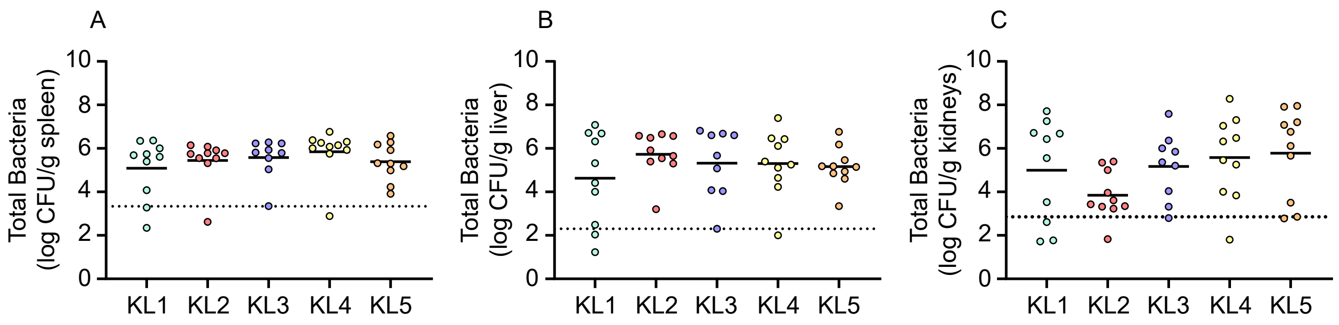


**Supplemental Figure 2. Bacterial burdens from TVI competition infection experiments.** Total bacteria recovered from mixed competition infections were quantitated in the spleen (A), liver (B), and kidneys (C) following TVI bacteremia (24 h) in C57BL/6J mice (n=10). Solid lines represent the mean of log transformed values and the dotted lines represent the highest value among samples that were at or below the limit of detection. None of the strain mixtures had significantly different colonization densities within tested organs as assessed by one-way ANOVA with Tukey's multiple comparisons test.
