## Supplemental Figure 3 for "Infection characteristics among *Serratia marcescens* capsule lineages"

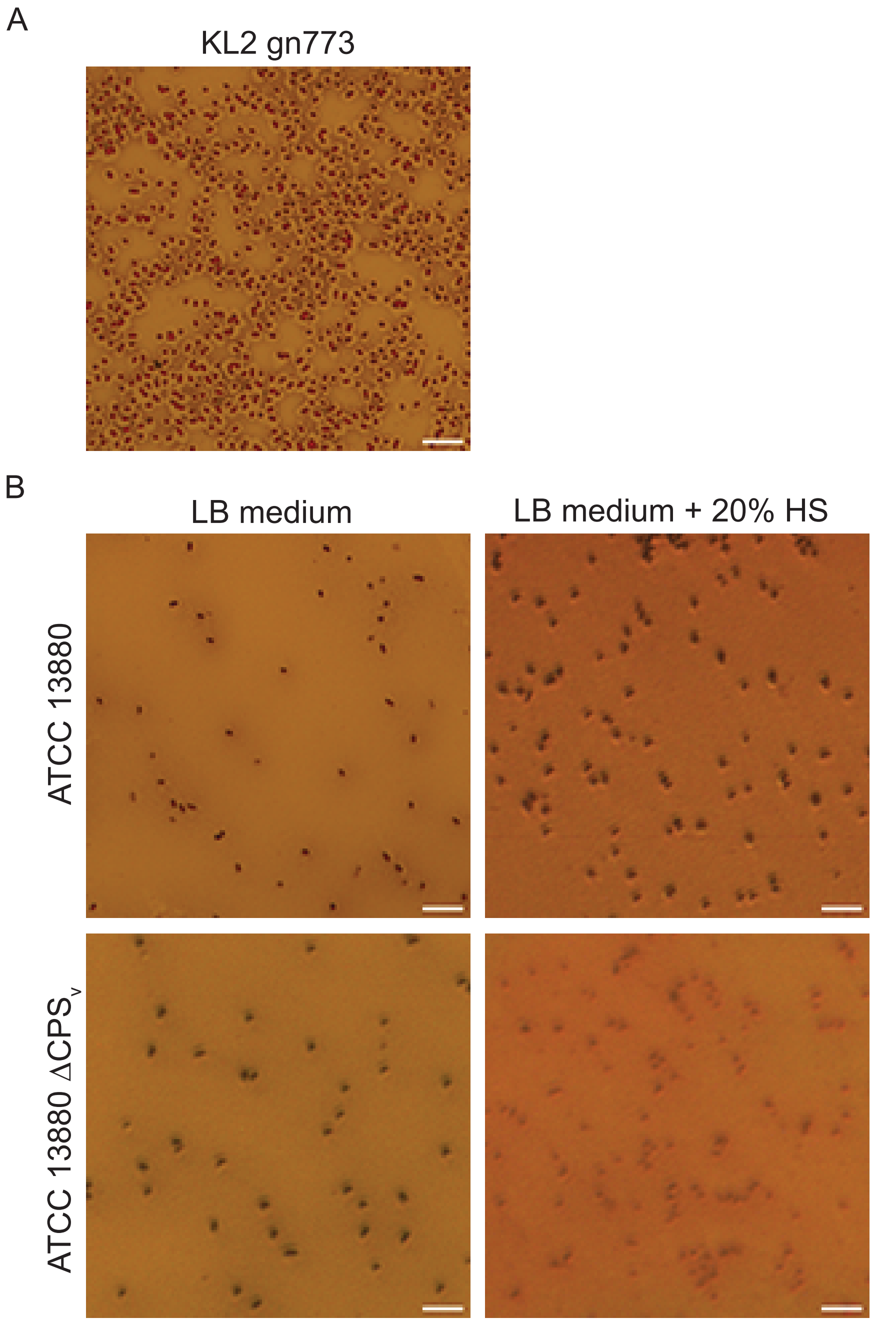


**Supplemental Figure 3. Serum exposure does not induce ATCC 13880 capsule synthesis.** A. A control culture of encapsulated strain KL2 gn773 was stained with Maneval's reagent, demonstrating a negative staining region surrounding gn773 cells indicative of capsule. B. ATCC 13880 and the $\Delta CPS_v$ derivative were passaged in increasing concentrations of human serum (HS) over the course of three days and subjected to Maneval stain after the final passage in 20% HS. No evidence of CPS synthesis was detected from either strain by this method. Scale bars are 5 μm.
