## Supplemental Figure 4 for "Infection characteristics among *Serratia marcescens* capsule lineages"

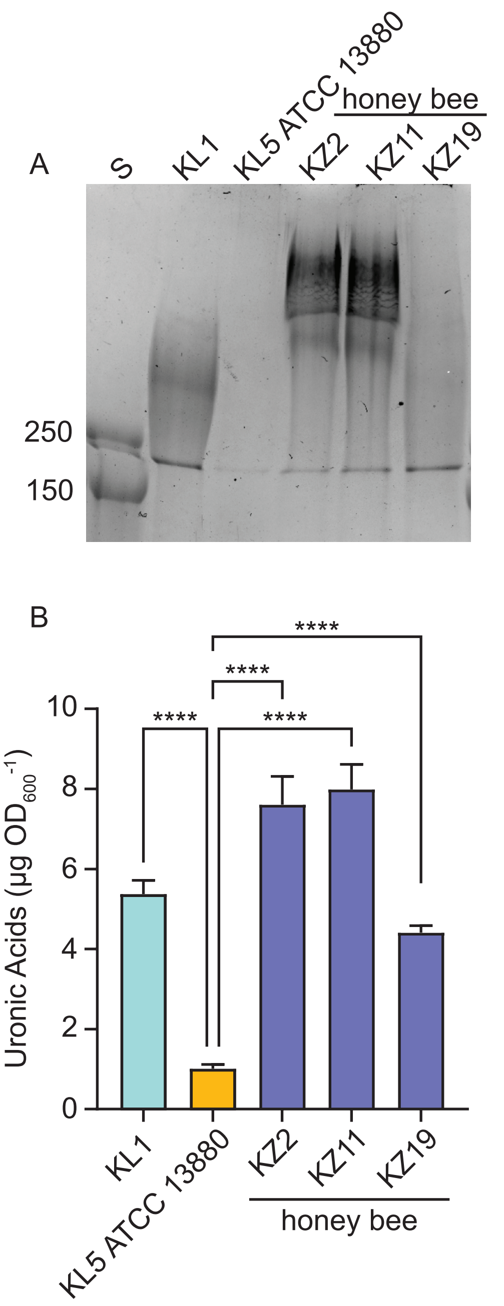


**Supplemental Figure 4. CPS production by non-clinical isolates.** A. Total polysaccharides were isolated from the indicated *S. marcescens* strains and separated by SDS-PAGE in comparison to protein standards (S) of known molecular weight (kDa). Gels were stained with alcian blue for visualization of CPS. B. Quantitation of extracellular uronic acids in *S. marcescens* strains. Statistical significance was assessed by one-way ANOVA with Dunnetts multiple comparisons test against ATCC 13880; ****, P < 0.0001.
