## Supplemental Figure 5 for "Infection characteristics among *Serratia marcescens* capsule lineages"

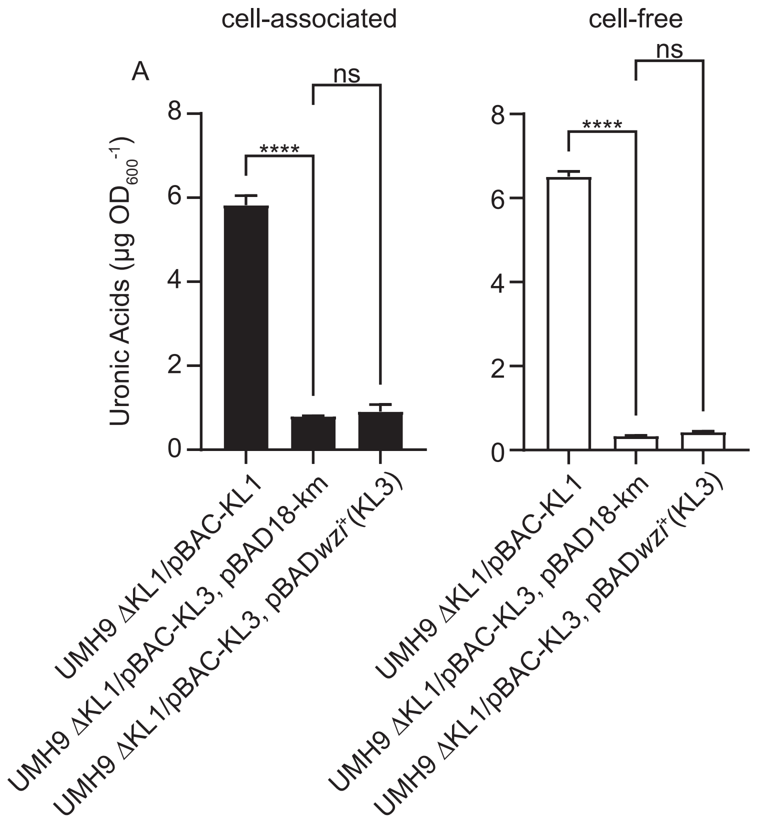


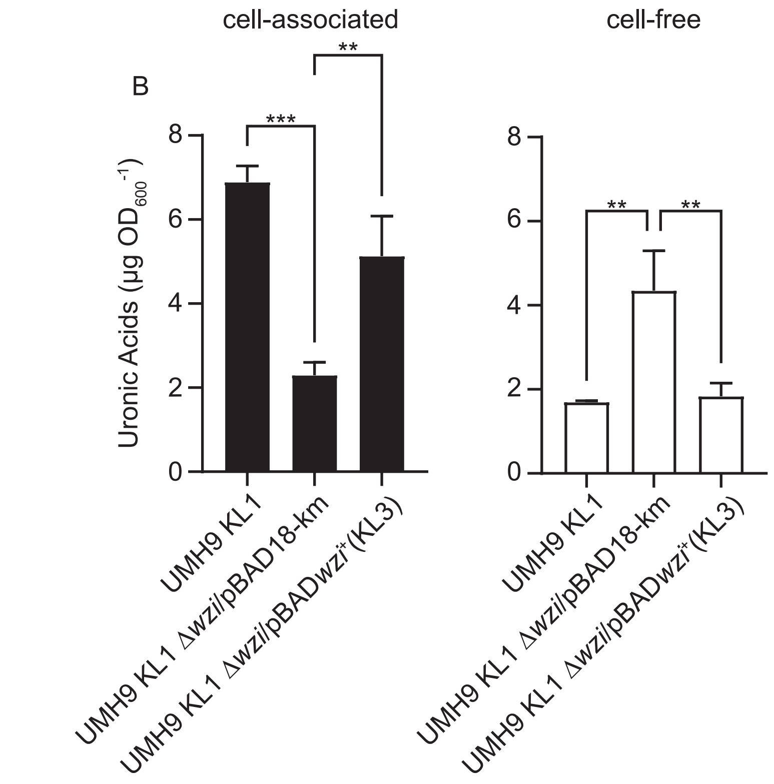


**Supplemental Figure 5. Genetic complementation of CPS cell-association by heterologous *wzi* gene expression.** A. Expression of the UMH7 KL3 *wzi* gene in the UMH9 ΔKL1/pBAC-KL1 background. B. Expression of the UMH7 KL3 *wzi* gene in the UMH9 KL1 Δ*wzi* strain. Uronic acids were quantitated from pelleted bacterial cells (cell-associated) or filter-sterilized culture supernatants (cell-free) in comparison with a glucuronic acid standard curve. Statistical signficance was assessed by one-way ANOVA with Dunnett's multiple comparisons test relative to strains harboring the vector control plasmid pBAD18-km: ns, not significant; **, Adj. P <0.01; ***, Adj. P <0.001; ****, Adj. P <0.0001.
