## Supplemental Figure 6 for "Infection characteristics among *Serratia marcescens* capsule lineages"

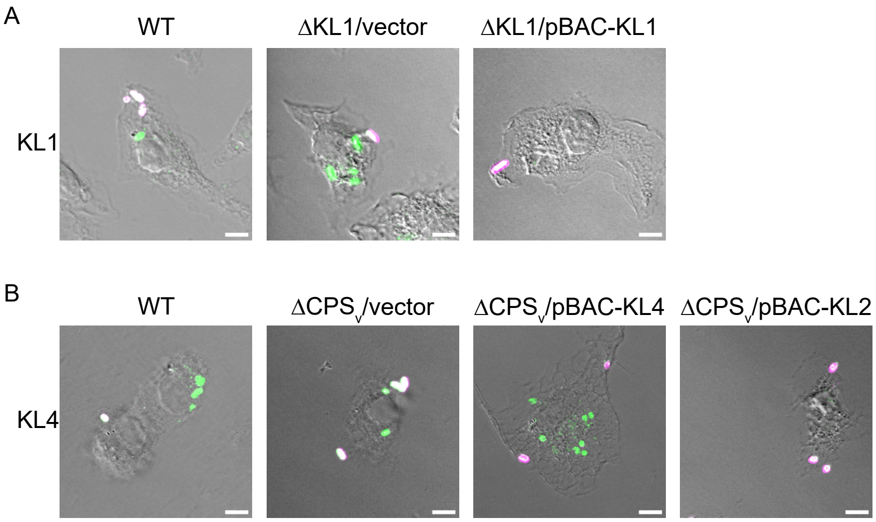


**Supplemental Figure 6. KL1 and KL2 CPS limit macrophage phagocytosis of *S. marcescens*.** Representative images of wild-type (WT) and capsule mutant derivatives of KL1 (A) and KL4 (B) in association with BMDM after 60 min infection. Extracellular bacteria were labeled with an AlexaFluor-647 conjugated secondary antibody, BMDM were then permeabilized and all bacteria were exposed to an AlexaFluor-488 conjugated secondary antibody. Extracellular bacteria fluoresce in both channels and appear white in the composite images while intracellular bacteria appear green. Scale bars are 5 µm. The WT KL1 image in panel A is the same as shown in Figure 10.
