## Supplemental Figure 7 for "Infection characteristics among *Serratia marcescens* capsule lineages"

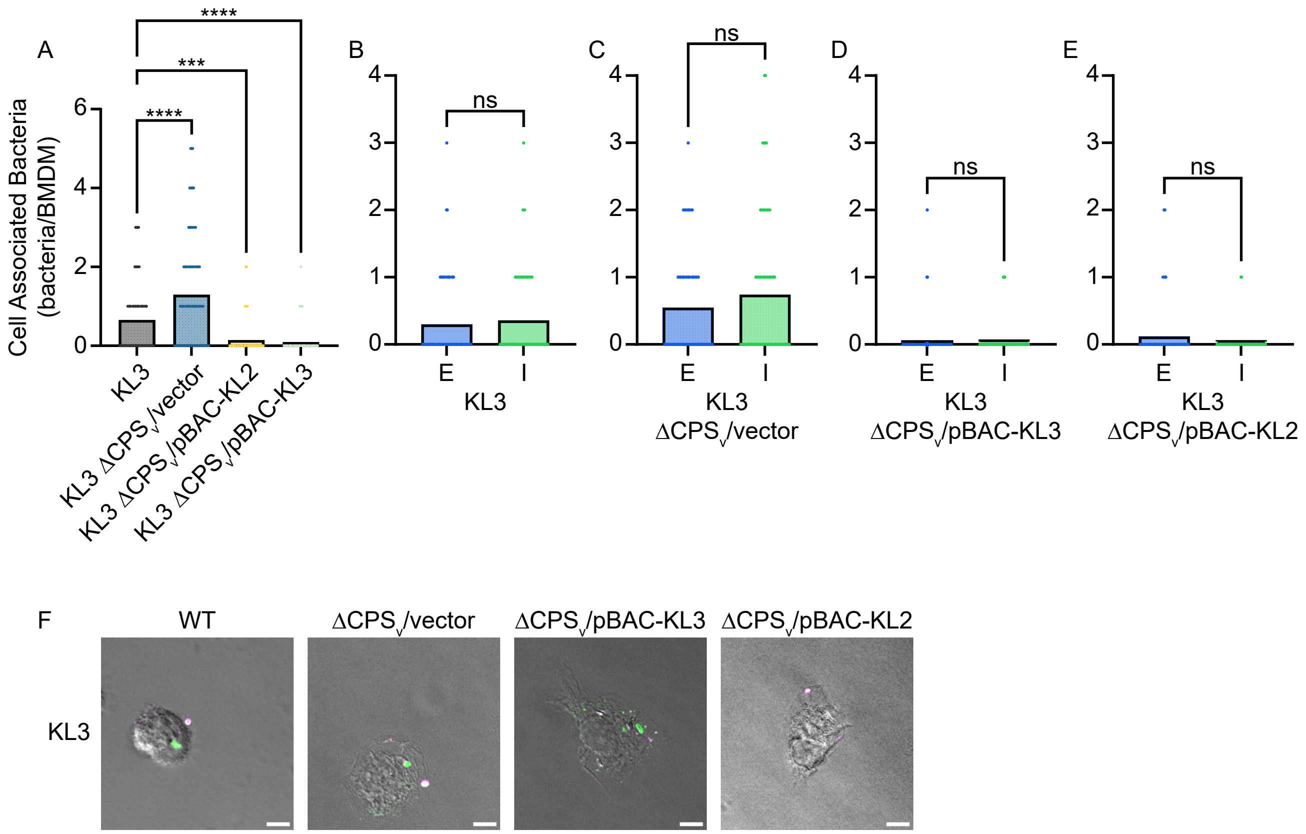


**Supplemental Figure 7. KL3 CPS does not alter localization of *S. marcescens* in association with BMDM**. A. Total BMDM-associated (intracellular + extracellular) bacteria were quantitated after 60 min co-incubation via differential immunofluorescence and microscopy. Points represent the number of bacteria associated with individual cells and statistical significance was assessed by one-way ANOVA with Dunnett's multiple comparisons test against the wild-type KL3 control strain. Adj. P: ***, <0.001; ****, <0.0001. B-E. Extracellular (E) and intracellular (I) bacteria were enumerated on a per cell basis for wild-type KL3 and derivatives of the $\Delta CPS_v$ KL3 mutant strain. Differences between intracellular and extracellular BMDM-associated bacteria were determined to be non-significant (ns) by unpaired t-test. F. Representative composite images from KL3 wild-type (WT) and capsule mutant strain BMDM infections. Extracellular bacteria were first labeled with an AlexaFluor-647 conjugated secondary antibody, BMDM were then permeabilized and all bacteria were exposed to an AlexaFluor-488 conjugated secondary antibody. Extracellular bacteria fluoresce in both channels and appear white while intracellular bacteria appear green. Scale bars are 5 μm.
